# Single cell multi-omic mapping of subclonal architecture and pathway phenotype in primary gastric and metastatic colon cancers

**DOI:** 10.1101/2022.07.03.498616

**Authors:** Xiangqi Bai, Billy Lau, Susan M. Grimes, Anuja Sathe, Hanlee P. Ji

**Affiliations:** Division of Oncology, Department of Medicine, Stanford University School of Medicine, Stanford, CA, United States; Department of Electrical Engineering, Stanford University, Palo Alto, CA, United States

**Keywords:** Single cell DNA sequencing, RNA sequencing, gastric and colorectal cancer, subclone architecture, multi-omics integration, biological pathways

## Abstract

Single cell genomics provides a high-resolution profile of intratumoral heterogeneity and subclonal structure in primary and metastatic tumors. Notably, metastases and therapeutic resistant tumors often originate from distinct subclones. These distinct cellular populations are an important contributor to adaptation and resistance to ongoing therapy. Single cell DNA-sequencing (**scDNA-seq**) defines subclones but does not provide biological information about cell types. Single cell RNA-sequencing (**scRNA-seq**) provides biological information but is less useful for identifying different subclones. The integrated scDNA-seq and scRNA-seq data from the same tumor cell population provides both subclone structure and biology. To understand the cellular genomic variation of different subclones in primary and metastatic cancers, we developed an approach to integrate multi-omics data from both types of single cells. This joint data set represented thousands of normal and tumor cells derived from a set of primary gastric and metastatic colorectal cancers. The extensive cellular sampling provided robust characterization of the subclonal architecture of gastric and colorectal cancers. We reconstructed the subclonal architecture using the cells in G0/G1 phase. The scDNA-seq provided a ground truth for copy number-based subclones. From the scRNA-seq data, the epithelial cells in G0/G1 were identified and assigned to specific subclones by a correlation algorithm based on gene dosage. The inferred CNV profiles from the scRNA-seq epithelial cells were assigned subclones identified from the scDNA-seq. Afterward, we determined the biological pathway activities of specific clones. Overall, integrative multi-omics analysis of single-cell datasets is more informative than any individual genomic modality, provides deep insights into the intratumoral heterogeneity, and reveals subclonal biology.

## BACKGROUND

Cellular genomic heterogeneity is a fundamental feature of cancer. Subclonal cellular populations are present within a tumor and have different genetic and biological properties. Complex subclonal architecture is present in primary tumors, metastases, patient-derived xenografts, and even cancer cell lines [1–3]. Notably, the dominant subclones of resistant tumors often originate from minor subclones in the primary tumors, as has been demonstrated in gastric cancer [4]. Single-cell genomics has provided a way to define subclonal architecture. Single-cell DNA sequencing (**scDNA-seq**) approaches include low coverage whole genome sequencing (**WGS**) for identifying somatic copy number variation (**CNVs**) or targeted sequencing for detecting cancer mutations [2, 3, 5–8]. The genomic CNV features define subclonal DNA characteristics in a tumor. Moreover, cancer CNVs are genomic indicators of tumor evolution including adaptation to ongoing therapy [9]. However, scDNA-seq does not identify cell types or other cellular phenotypic features. Other single-cell methods provide features relevant to phenotype and biology. Namely, single-cell RNA sequencing (**scRNA-seq**) delineates the gene expression of cancer cells. While scRNA-seq does not provide a DNA-based genetic definition of subclones, one important feature to infer the underlying genome structure of intratumoral heterogeneity [10]. This information denotes the malignant characteristics underlying clonal expansion, proliferation, metastasis, and treatment response in cancer [1, 11]. By combining the two single cell assays, one identifies underlying genomic alterations among the cells in the sample, characterizes subclonal cellular diversity; and reveal biological pathways in specific cellular populations [3, 12–14]. Overall, the integration of both single-cell DNA and RNA assays from the same tumor provides a highly informative measurement of tumor biology [1, 11].

There are several algorithms for multi-omics integration of scDNA-seq and scRNA-seq. Generally, these algorithms use map-to-reference inference methods such as clonealign [15] and Seurat [16]. The scRNA-seq data is integrated with the DNA-based clonal features. However, these methods do not provide strong correspondence between paired scDNA and scRNA of primary tumors. This issue results of the high intratumoral heterogeneity found in primary tumors and variable performance of CNV calling from the single-cell platforms.

We developed a comprehensive workflow for single-cell copy number analysis, then performed an integrated analysis on these two single-cell platforms. This study used single cell suspensions derived from primary or metastatic tumor biopsies. Using the same cell suspension from a given cancer, we conducted scDNA-seq and scRNA-seq assays on hundreds to thousands of cells. The extensive cellular sampling provided robust single cell characterization of both DNA and RNA-based features from the same tumor. The scDNA-seq analysis process includes cell quality control, cellular component identification, feature selection, and subclonal reconstruction. Each subclones had a distinct set of CNVs, typically involving large chromosome segments. Subsequently, we analyzed the corresponding scRNA-seq gene expression data to determine cell types and gene expression characteristics. Using both single-cell DNA and RNA results, we utilized a correlation algorithm to assign cells from the scRNA-seq data to specific subclones of scDNA-seq based on the gene dosage effects [17–19]. Finally, we applied the GSVA to identify differential pathway activities in each subclone.

We used this single-cell multi-omics approach to define the subclonal architecture and clonal phenotype from a series of gastrointestinal cancers in the stomach and colon. We compared the matched primary colorectal cancers (**CRCs**) and their metastasis. Notably, this study provides a high-resolution subclonal architecture for these digestive tract cancers and identified subclone-specific biological pathway information.

## MATERIALS AND METHODS

### Samples

This study was conducted in compliance with the Helsinki Declaration. All patients were enrolled according to a study protocol approved by the Stanford University School of Medicine Institutional Review Board (IRB-44036). Informed consent was obtained from all patients. Tissues were obtained from the Stanford Cancer Institute Tissue bank. We also had matched adjacent normal tissue for a subset of the samples. Ascertainment of samples was based on the availability and cellularity of matched normal tumor samples that occurred from 2015 to 2019. All the specimens underwent histopathology review with hematoxylin and eosin-stained tissue sections.

After surgical resection, the tumor samples were stored in RPMI medium on ice for less than 1 hour. The sample was then macrodissected and dissociated into a cellular suspension by the gentleMACS Octo Dissociator using the human tumor dissociation kit as per manufacturer’s recommendations and the 37C_h_TDK_3 program (Miltenyi Biotec, Bergisch Gladbach, Germany). The suspension was used for scRNA-seq. Single cell DNA-Seq was performed after thawing cryopreserved dissociated cells stored in liquid nitrogen in 90% FBS-10% DMSO freezing medium.

### Single cell DNA-Seq library preparation and sequencing

Single-cell DNA libraries were generated using a high-throughput, droplet-based reagent delivery system using a two-stage microfluidic procedure (10X Genomics Inc., Pleasanton, CA). First, cells were encapsulated in a hydrogel matrix and treated to lyse and unpackage DNA. Second, a gel bead was functionalized with copies of a unique droplet-identifying barcode (sampled from a pool of ~737,000) and co-encapsulated with the hydrogel cell bead in a second microfluidic stage to separately index the genomic DNA of each individual cell.

In the first microfluidic chip, cell beads were generated by partitioning approximately 10,000 cells of each sample in a hydrogel matrix. A cell suspension is combined with an activation reagent, hydrogel precursors, paramagnetic particles, and loaded into one inlet well. In the other two inlet wells, a polymer reagent and partitioning oil were added. To ensure a low multiplet rate, cells were delivered at a dilution such that the majority of cell beads contain either a single cell or no cell. Once generated, the emulsion was immediately transferred into a PCR strip tube and incubated with orbital shaking at 1000 rpm overnight. The incubation yields polymerized magnetic cell beads for subsequent steps.

Encapsulated cells were processed by the addition of lysis and protein digestion reagents to yield accessible DNA for whole-genome amplification. After the lysis, cell beads were washed by magnetic capture, concentrated by reduction of liquid volume, and buffer exchanged with the addition of 1X PBS buffer. Cell beads were then denatured by NaOH, neutralized with Tris, and diluted in storage buffer. Finally, aggregates of cell beads were removed by filtration through a Flowmi strainer before a volume normalization procedure to set the cell bead concentration. To amplify and barcode gDNA, the emulsion was then incubated at 30°C for 3 hours, 16°C for 5 hours, and finally heat inactivated at 65°C for 10 minutes before a 4°C hold step. This two-step isothermal incubation yielded genomic DNA fragments tagged with an Illumina read 1 adapter followed by a partition-identifying 16bp barcode sequence.

The emulsion was broken and purified per the manufacturer’s protocol. Conventional end-repair and a-tailing of the amplified library was performed, after which a single-end sequencing adapter containing the Illumina read 2 priming site was ligated. PCR was performed using the Illumina P5 sequence and a sample barcode with the following conditions: 98°C for 45 seconds, followed by 12-14 cycles (dependent on cell loading) of 98°C for 20 seconds, 54°C for 30 seconds, and 72°C for 30 seconds. A final incubation step at 72°C was performed for 1 minute before holding at 4°C. Libraries were purified with SPRIselect beads (Beckman Coulter, Brea, CA) and size-selected to ~550bp. Finally, sequencing libraries were quantified by qPCR before sequencing on the Illumina platform using NovaSeq S2 chemistry with 2×100 paired-end reads.

### Single cell DNA-Seq data processing and CNV calling

Sequencing data was processed with the cellranger-dna pipeline (6002.16.0), which automates sample demultiplexing, read alignment, CNV calling, and report generation. Paired-end FASTQ files and a reference genome (GRCh38) are used as input. The cellranger-dna pipeline includes preprocessing and single cell copy number calling parts. It outputs CNV calls and read counts in 20kb bins across whole genome for individual cells as genomic bin-by-cell matrices. In the preprocessing stage, the first 16 base pairs of read 1 are compared to a whitelist of all possible droplet barcodes (totaling ~737,000). All the observed droplet barcodes were tested for the presence of a cell by using mapped read abundances to the human genome. Reads were aligned to GRCh38 using bwa-mem. Each read in the bam file was annotated with a cellular barcode tag ‘CB’. High confidence mapped reads were counted across the genome in 20kb non-overlapping windows. GC bias correction, modelled as a polynomial of degree 2 with fixed intercept, was applied. Copy number calls were generated through modeling binned read abundances to a Poisson distribution with the copy number, GC bias, and a scaling factor as parameters. Candidate breakpoints were estimated by applying a log-likelihood ratio statistic against fluctuations in read coverage over neighboring genomic bins. These breakpoints were refined and reported as a set of non-overlapping segments across the genome. The copy numbers were scaled to integer-level ploidies. Copy number calls for non-mappable regions were imputed with neighboring copy number calls in confidently mapped regions, provided that the copy number on both sides of a non-mappable region were the same and the region was < 500 kb.

### Matrix format of single-cell CNV calling

The single-cell CNV calling output of cellranger-dna pipeline is the “node_cnv_calls.bed”. We converted the single-cell CNV calls into a matrix format with consistent genome bins to make it suitable for dimensional reduction and clustering approaches. This is similar to the scRNA-seq analysis. Specifically, the entire genome segments of cells in a “node_cnv_calls.bed” file were derived and mapped to unified evenly 20kb bins on the reference genome (GRCh38). A bin-by-cell scDNA-seq integer CNV matrix was generated for each sample with a total of 154423 non-overlapping 20kb genome as row features.

We merged the 50 neighbored 20kb bins into one 1Mb-level segment in each chromosome. The copy number variants per 1Mb segment was calculated as the median of CNV across 50 neighborhood bins. Finally, the high dimensional 154423 bins were aggregated to 3103 1Mb-level segments on the WGS. The single-cell CNV matrix with 3103 genomic segments/features was used for downstream analysis.

### Quality control for scDNA-seq data

For the high-throughput single-cell DNA sequencing technology, a substantial proportion of cells are deemed as noisy cells due to underlying various factors: (i) input cells or nuclei suspension’s DNA degraded; (ii) technical noise during workflow. These two factors may cause large dropout proportion of single cells. Here, we identified the single-cell DNA dropout by computing the ratio of the number of segments (copy number = 0) based on a total of 3103. We detected technical noise cells when setting a uniform threshold 0.1 for the dropout ration across current samples (**Additional File 1: Figure S1**).

Other than the experimental technical noise, there were still part cells labeled as cellranger-dna noise as shown in the “per_cell_summary_metrics.csv” file. Those high noise cells were labelled with either “High DIMAPD” or “Low ploidy confidence” as determined through a cellranger statistical model. We noted that a part of cellranger-dna noise cells belongs to actively replicating cells in S phase. We identified the S phase cells if their CNVs were larger than 2.5.

### Normal cell type identification of tumor scDNA-seq data

The tumor tissues were partly composed of normal diploid cells such as immune and stromal cells in the microenvironment. To identify the normal cells in the tumor sample, we merged the 2888 segments without the X and Y genomic segments into 22 autosomes, and then calculated the CNV mean of all segments from the same chromosome. This summation enabled one to identify large chromosomal alterations. We defined normal cells in which all the chromosomal-CNVs on the 22 autosomes had a copy number value less than 2.5.

### Cell cycle assignment of scDNA-seq tumor cells

To distinguish G0/G1 cells and S phase cells, we computed the Euclidean distance among the cells for a given sample. This involved considering those cells with a copy number equal to 2 across the 3103 1Mb-level segments. Replicating cells showed higher dissimilarity distance since because the CNV values generally double compared to when they are in G0/G1 phase. We utilized an Expectation-Maximization (**EM**) algorithm to fit a 2-component normal mixture distribution for obtained Euclidean distance vector (**Additional File 1: Figure S2 A**). The replicating cells were automatically assigned to the normal distribution with the larger mean, otherwise the rest cells located at normal distribution with small mean were assigned to G0/G1 phase. Finally, we separated G0/G1 phase cells and S phase cells. Some of the S phase cells had a degree of noise that would potentially exclude them. We classified these replicating cells if their CNVs were greater than 2.5.

### Genomic instability analysis in focal events

We quantitatively measured the level of genomic instability for a given tumor. This analysis involved filtering out all noisy cells described previously. For each sample, we estimated all consecutive genome regions with genomic instability (copy number ≠ 2) in “node_cnv_calls.bed”. Subsequently, we counted the presence of a CNV across genome intervals of different sizes as follows: ≤ *20kb*; *20kb*~*100kb*; *100kb*~*500kb*; *500kb*~*1Mb*; *1Mb*~*5Mb*; *5Mb*~*10Mb*, ≥ *10Mb*. We also calculated the proportion of amplification and deletion events in total genomic instability for all samples.

Chromosomal instability (**CIN**) is a source of genomic heterogeneity in a population of gastric tumor cells. The extent of chromosomal CIN on a single cell basis was quantified through the “node_cnv_calls.bed” file. This is one of the outputs from cellranger-dna pipeline. It contained copy number calls on mappable genomic regions for every single cell with imputation. For each single cell, we counted the frequency of genomic segments with aneuploidy (CNV≠2), then calculated the percentage of CIN in total number of genome regions on the WGS.

### Integration of single-cell CNVs with TCGA CNVs

As a comparison to existing cohorts, we compared the aggregate CNV structure on chromosomal arms of primary gastric and colorectal cancer samples with known chromosomal patterns provided from public cancer genome data sources. We downloaded the bulk DNA-seq data of the TCGA STAD samples (https://www.cbioportal.org/study/summary?id=stad_tcga_pan_can_atlas_2018, https://www.cbioportal.org/study/summary?id=stad_tcga). The putative copy number alterations profile from total 411 STAD samples were processed by calculating the mean of CNV across the genome segments at the same chromosome arm. We determined the frequency of amplifications and deletions on arm levels can be conducted for bulk data. Meanwhile, we aggerated single-cell CNV files of four primary gastric tumor samples but excluding all noisy cells. Then we computed the frequency of non-noisy single cells with CNV gains or deletions on each arm. Finally, we selected the top 10 frequent chromosomal arms with either amplification or deletion and compared the common results between TCGA samples and single-cell datasets.

### Subclonal reconstruction with scDNA-seq

The noisy cells, normal cells and replicating tumor cells were eliminated prior to Seurat processing. Seurat (v.3.1.2) was applied to the G0/G1 tumor cells single-cell CNV matrix. This output was used to construct the subclonal structure through unsupervised clustering. There was a similar pre-processing procedure with scRNA-seq analysis that included cell filtering and feature selection. We developed an alternative statistical approach to select most informative genomic segments with high variance. This alternative was essential to reduce the high dimension and improve the clustering performance. The detailed steps include the following: (i) We fitted a mixture normal distribution to the inter-cell copy number variances of all 3103 segments by EM algorithm (**Additional File 1: Figure S2 B**). (ii) We iteratively selected the most informative segments with the highest variance which were located at the larger mean normal distribution, until at least 20% of the total segments were included. The selected informative copy number segments were used as the features in scDNA matrix input to Seurat.

We utilized the ‘CreateSeuratObject’ function to the processed input matrix and did not set any filtering parameters. We applied ‘RunPCA’ function to the raw CNV matrix without any normalization and scale. The first 20 principal components (**PCs**) were used to “RunUMAP” function. We implemented the Louvain algorithm to detect the optimal clustering on a shared nearest neighbor (**SNN**) graph which was constructed on the two-dimensional UMAP embedded space. Noted that, the resolution parameter in ‘FindClusters’ function affects the number of clusters. We set the resolution parameter relying on the number of cells as shown below.

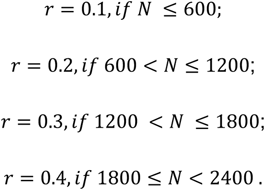

Where *r* means the “resolution” parameter in “FindClusters”, the *N* means the cell number. Finally, Seurat defined the major clusters of G0/G1 tumor cells. Subclones consisting of several cells were difficult to detected through automatically clustering algorithm. Thus, we used inspection-based K-means clustering to separate the minor subclones with only a small number of cells.

### Phylogeny tree construction

For the subclonal phylogeny, the subclonal CNVs were calculated as the integer average of CNV values of single cells in that subclone. The “dist” function on stats (v.3.6.2) package was used to calculate the Euclidean distance matrix of subclonal CNVs vectors. We performed hierarchical clustering for the distance matrix by using ‘hclust’ function with default parameters. The R package ape (v.5.4) [20] was then applied to construct the phylogenetic tree of subclones utilizing “plot.phylo” function. To visualize the CNV changes among subclones, we used ComplexHeatmap (v.2.5.4) [21] package – it provided a single-cell matrix with 1Mb-level segments on autosomes and cells were labeled by subclone labels. Meanwhile, the cells per subclone are sorted by ascending order of distance to standard normal cell (copy number = 2) in the heatmap. The subclones were also ordered according to the phylogenetic tree.

### Single cell RNA-seq

Cellranger software suite 3.0.2 was used to process the scRNA data. This included sample demultiplexing, barcode processing and single cell 3′ gene counting. The cDNA sequence from read 2 was aligned to the GRCh38 human reference genome. The reference GTF contained 33,694 entries, including 20,237 genes, 2337 pseudogenes and 5560 antisense (non-coding DNA). Cellranger provided a gene-by-cell matrix, containing the read count distribution of each gene for each cell.

The cellranger program provide the raw gene expression count matrix per sample. The Seurat package (v.3.1.2) was used to perform cell and gene quality control. Filters included the following: (i) genes expressed in less than three cells; (ii) cells expressing fewer than 200 genes and over 5000 genes; (iii) cells expressing at least 30% mitochondrial transcripts. The cell cycle scores for the G2M and S phases were determined using the “CellCycleScoring” function. Doublet cells with an outlier range of UMI were computationally detected by DoubletFinder (v.2.0.1) [22]. We excluded the doublets from downstream analysis. Following above quality control steps, high quality cells were used for downstream analysis.

We eliminated samples with a low number of expressed genes number across cells (i.e., P5847_normal, P6342_normal, P6461_metastasis, P6593_tumor). To ensure the most of poor-quality cells were filtered out, we adjusted the filtering parameters during data quality control after manually reviewing the results (**see Additional File 2: Table S6**).

For each patient, we used the “merge” function in ‘SeuratObject’ (v.4.0.4) package to aggerate the paired samples’ objects. Specifically, the normal and tumor samples were merged for four primary gastric tumors. A similar step was used for the colorectal cancer single cell data. We utilized “SCTransform” function to normalize the UMI count matrix and select the top 3000 variables genes. The normalized matrix with high variable genes were used as input to “RunPCA” function. We constructed a shared nearest neighbor graph based on the first 20 principal components (**PCs**) – this step used the “FindNeighbors” function. Finally, the ‘FindClusters’ function with the resolution parameter 0.8 was implemented to identify the optimal cell clusters on the constructed SNN graph.

### Epithelial cell detection, re-clustering and visualization

To identify the epithelial cells from each sample, we compared the average epithelial module score (**MS**) and non-epithelial MS in each cell cluster. This score was used to assign cells as being epithelial versus other cell types. We employed the following steps: (i) Calculating the epithelial MS of individual cells by adding the epithelial cell marker list into “AddModuleScore” function, in which the gene list was searched on public literature as (*MUC5AC, PGC, EPCAM, MUC2, TFF1*). The same procedure was used to compute the non-epithelial MS for each cell, in which non-epithelial markers mainly composed by other cell lineages biomarkers, such as fibroblasts (*ACTA2, DCN, SPARC, THY1*), endothelium (*VWF, PECAM1*), dendritic (*CD83*), macrophage (*CD14, FCGR3A*), immune (*PTRPC*), B plasma (*CD19, CD79, MS4A1, IGHA, IGHC, IGLC*) and T cells (*CD3D, TRAC, TRAB, CD8A, NKG7, GNLY, IL2RA*). (ii) The minimum positive average non-epithelial MS in clusters which average epithelial MS were less than 0 was chosen as the threshold to cut off the epithelial cell clusters. For epithelial cell assignment we identified the clusters with epithelial MS greater than 0.

We used Seurat to re-cluster the epithelial cells of aggregated object at a higher resolution. The main procedure was similar with the former analysis pipeline. The “SCTransform” with the parameter “nCount_RNA” was applied to regress out the variation effects in sequencing depth, and including normalization, scaling and variable feature selection. The first 100 principal components identified by ‘RunPCA’ were used for unsupervised clustering with a resolution set to 0.8. UMAP was conducted on the top 100 PCs as further dimensional reduction for visualization.

### Assignment of scRNA-seq cells to scDNA-seq subclones

With the paired scDNA-seq and scRNA-seq data from the same tumor, we integrated the two modalities. This analysis constructed the corresponding alignment between transcriptomic subclusters and genomic subclones. The subclone architecture based on scDNA-seq was regarded as the ground truth. We utilized gene dosage relationship as the basis for the correlation method to assigning scRNA-seq cells to subclones. This method consists of the following steps: (i) normalize the expression matrix using “LogNormalize” function in Seurat package with the log-transform being applied to the gene expression of each cell after scaling by the total expression multiples 10000; (ii) select the genes on the amplified and deleted chromosomes contributed to the subclones; (iii) compute the Spearman correlation *ρ_nc_* for the scaled expression of the selected genes for cell *n* the subclone *c* average gene-level copy number profile; (vi) assign cells to the subclone with highest correlation, *z_n_* = *arg* max *ρ_nc_* We determined the correlative gene expression related to gene dosage by the copy number changes.

### Copy number analysis of scRNA-seq data

We applied inferCNV [23] (v.1.2.2) (https://github.com/broadinstitute/inferCNV) to estimate the somatic large-scale chromosomal copy number alternations from scRNA-seq profile of epithelial cells. The raw count matrix with genes by cells was used as input. We also included cell cluster and gene annotation files. The normal epithelial cells originated were used an internal baseline reference group (**Additional File 2: Table S8**). The metastatic tumors did not have matching normal tissue. Therefore, we set the reference cells as “NULL”. Moreover, the parameters were set as “cutoff=0.1”, “denoise=T” and “HMM=T” in inferCNV to enable the CNV predictions accuracy. Finally, the relative large-scale CNVs and aneuploidy status of tumor subclones were inferred through comparing with normal reference group. The inferCNV i6 HMM results predict CNV levels of gene expression profile. We determined which of the inferred CNVs were consistent with this scDNA-seq CNV. Then we plotted a heatmap to visualize the inferred CNV changes among the different subclones defined from the scRNA-seq.

### Statistical analysis

We used Spearman correlation and root mean square error (**RMSE**) to evaluate the integration performance. The inferred CNV matrix with genes of scRNA-seq was annotated as *X_n×g_*. The CNV matrix of scDNA-seq subclones with the common genes *g* was represented as *S_c×g_*. We aggregated the CNVs on genes as the average CNVs on 22 autosomes for each subclone. Then the inferred CNV matrix *X_n×g_* was converted as 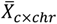, and the subclones’ CNV matrix *S_c×g_* as 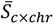. The similarity between inferred CNV pattern of scRNA-seq clonal assignment 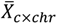 with ground truth 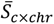 was calculated by Spearman correlation, as 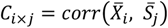, *i* =1,….,*c*; *j* = 1,…,*c*. We applied the RMSE metric to evaluate the residuals from the scRNA-seq estimated subclones to ground truth on genes resolution. In particular, the RMSE of the *i*-*th* subclone was calculated as 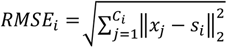, where the *x_j_* is the row of *X_n×g_*, *s_i_* is the *i-th* subclone of *S_c×g_*, *C_i_* is the set of cells assigned into the *i-th* subclone. The final RMSE was the average of RMSEs of all subclones as 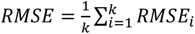, where *k* represents the number of subclones.

### Pathway analysis

We conducted a pathway enrichment analysis based on cancer hallmark signatures. This method defined the biological properties of epithelial cell clusters. We downloaded the hallmark gene set with 50 pathways from MSigDB (v.6.2) [24, 25] and imported them to ‘getGmt’ function in GSEABase (v1.46.0) [26] as a uniform format for downstream analysis. We used the Gene Set Variation Analysis (**GSVA**) (v.1.32.0) [25] to calculate enrichment scores for each pathway gene set of single cells using raw count matrix with 3000 high variable features as input. The Gaussian/Poisson kernel was set to estimate the non-parametric probability distribution of expression levels through parameter “kcdf = Gaussian” in ‘gsva’ function as well as the parameters “mx.diff=TRUE” and “min.sz=10”. We compared the enrichment scores among the clusters by performing ANOVA test. We selected the statistically significant pathways for each cluster if the false discover rate adjusted P-value < 0.05.

## RESULTS

### Approach overview

We performed scRNA-seq and scDNA-seq on gastrointestinal cancers which included four primary gastric adenocarcinomas and five metastatic CRCs with two having matched primary colon cancers (**Table 1, Additional File 2: Table S1**). The samples were from biopsies. The overall workflow of the single-cell multi-omics approach is shown in **Figure 1A**. For any given sample, we used the same cell suspension to prepare separate libraries for scDNA-seq and scRNA-seq assays. We had 13,255 scDNA-seq cells and 62,392 scRNA-seq cells from all tissue samples (**Additional File 2: Table S2, S3**).

**Figure 1.**
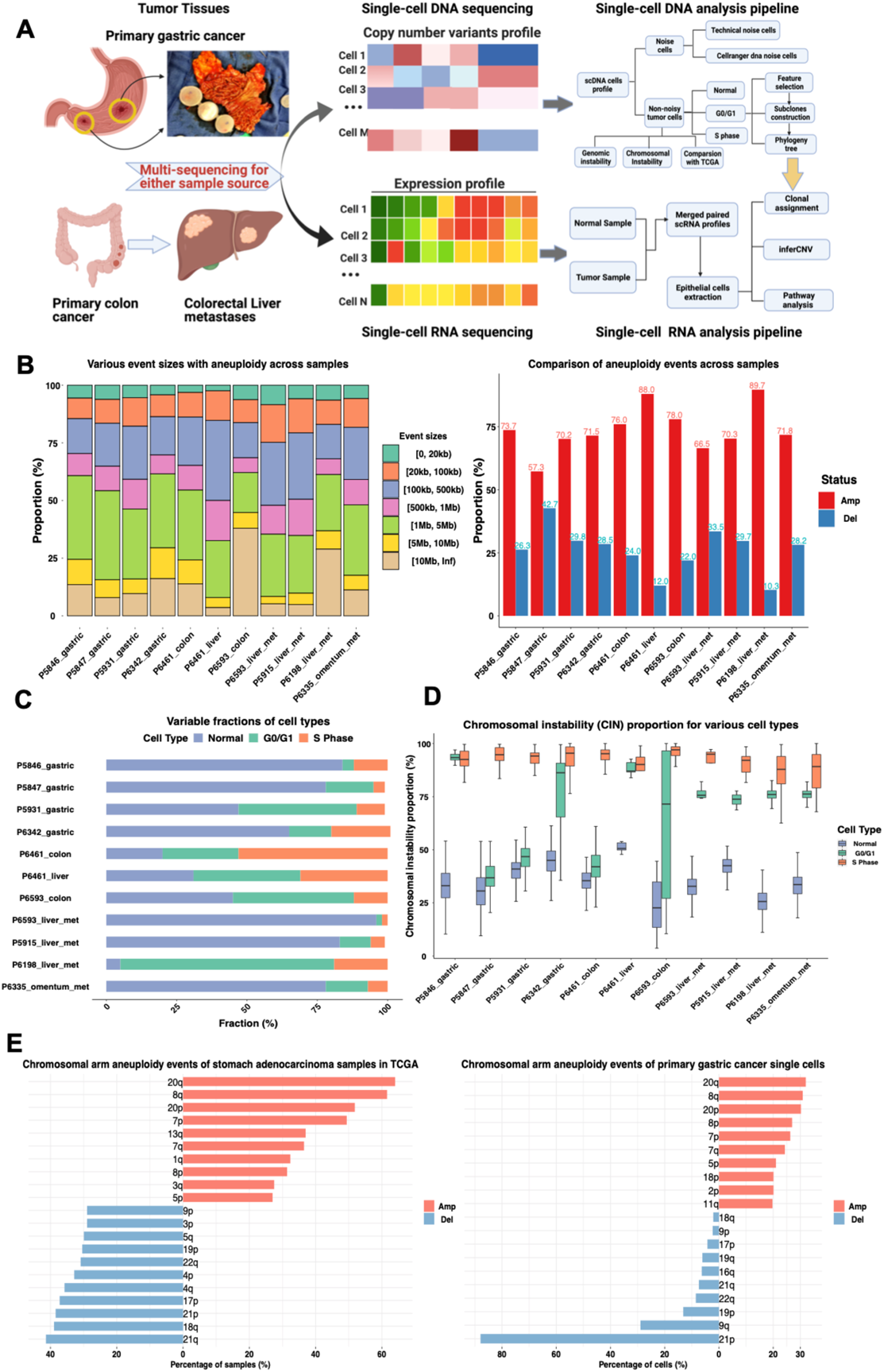
Study overview and evaluation of general genomic features. (A) Schematic diagram of study design and data analysis workflow. (B) The proportion of genomic instability sizes (left) and aneuploidy status (right) detected in single-cell DNA-seq samples. (C) The fraction of various cell types identified across samples. (D) Chromosomal instability (**CIN**) proportion of various cell types across samples. (E) The most frequent chromosomal arm occurring among the **TCGA STAD** tumors (left), and in measured primary gastric cancer single cells.

**Table 1.** Tumor samples.

| **Cancer type** | **Patient** | **Sample Type** | **Source** | **scDNA** | **scRNA** | **Paired normal tissue** |
| --- | --- | --- | --- | --- | --- | --- |
| **Gastric cancer** | **P5846** | Primary Tissue | Primary Tissue | Yes | Yes | scDNA & scRNA |
|  | **P5847** | Primary Tissue | Primary Tissue | Yes | Yes | scDNA & scRNA |
|  | **P5931** | Primary Tissue | Primary Tissue | Yes | Yes | scDNA & scRNA |
|  | **P6342** | Primary Tissue | Primary Tissue | Yes | Yes | scRNA |
| **Colorectal cancer** | **P5915** | CRC liver metastasis | Metastatic Tissue | Yes | Yes | Yes |
|  | **P6198** | CRC liver metastasis | Metastatic Tissue | Yes | Yes | Yes |
|  | **P6335** | CRC Omentum metastasis | Metastatic Tissue | Yes | Yes | scRNA |
|  | **P6461** | Matched colorectal primary / liver metastasis | Metastatic Tissue | Yes | Yes | scDNA (colon) |
|  | **P6593** | Matched colorectal primary / liver metastasis | Metastatic Tissue | Yes | Yes | None |

For scDNA-seq data, we filtered out the cells with low-quality data such as those with greater than 90% dropout of CNV segments. This reduced quality resulted from assay technical noise or cells with computationally inferred CNVs having a low statistical significance. After quality control filtering, the high-quality cell data was used to analyze tumors’ genomic instability and reconstruct the subclonal architecture.

Using the scDNA-seq data, we distinguished the normal cells with diploid content versus the tumor cell population. We established a threshold based on each cell’s average copy number value, removed the diploid cells, and identified the cells with aneuploidy. Based on the copy number value, cells were assigned to different cell cycle phases. We compensated for variation in the cellular DNA content by fitting a gaussian mixture model to the distance based on the ground truth CNV profile from normal diploid cells. Overall, the cells in G0/G1 provided the most consistent CNV calls.

We applied another statistical approach to determine the subclonal structure of G0/G1 aneuploidy cells. This method selects the most informative single-cell copy number features defining a subclone, embeds them into UMAP dimensional reduction, and uses a network-based Louvain clustering algorithm to find the distinct subclones on the integer copy number profile. Subsequently, we used a phylogenic tree algorithm to calculate the distance of consensus CNV profiles of each cluster. This analysis defined the individual subclones.

For scRNA-seq analysis, we extracted the epithelial cells from each sample. After applying the QC procedures to reduce noise and variance, we integrated the scRNA-seq results from the epithelial cells with scDNA-seq genomic subclone information. The algorithm uses a correlation assignment based on a gene dosage effect. The results demonstrated that the underlying subclonal structure for a given single-cell RNA-profile correlated with the ground truth subclonal architecture based on the scDNA-seq profiles. Based on the integration results, we identified the most significant biological pathways which differentiated each subclone’s biology.

### Single-cell analysis defines the global genomic instability of gastrointestinal cancers

We determined the extent of genomic instability among the different cancers [27–29]. We calculated the frequency of focal events of different segment sizes for each tumor. Among the primary gastric cancers, most CNVs ranged in segment size from 1Mb to 5Mb. This class of CNV segments contributed to approximately 30% of all genomic instability events per sample (**Figure 1B**). The number of CNV amplifications was higher than the deletion events regardless of the segment size (**Additional File 1: Figure S3**). In total, we observed an average 70% frequency for amplification events.

Among the CRCs, the metastases to the liver had the greatest degree of genome instability. The CNV amplification events per the metastatic colorectal tumors ranged from 66.6% to 89.7% (**Figure 1B**). The CNV segment lengths ranged from 100kb to 500kb. The P6593 colon cancer had a CNV segment greater than 10Mb in length.

### Evaluating the effect of cell cycle on copy number

Cells were assigned to different phases of the cell cycle. Normal cells were present in all samples and provided a useful control because their DNA content was consistently diploid [30]. The tumor cells were assigned to the G0/G1 or S [1]. Cells that were in the G0/G1 had a consistent CNV profile. In contrast, cells in the S phase had fluctuating CNV values because of DNA replication. We applied several steps to separate these cell cycle components. First, we assigned diploid cells to the ‘normal’ cluster (**Methods**). To distinguish tumor cells in G0/G1 and S phases, we used an Expectation-Maximization algorithm to estimate a 2-component normal mixture distribution for cell dissimilarities. Therefore, the cells originating from the higher mean normal distribution were automatically assigned to the S phase.

We determined that the percentage of the cells assigned to the normal cluster was between 47% to 84% across the four primary gastric cancer samples (**Figure 1C, Additional File 2: Table S4**). The proportion of S phase cells ranged from 4% in P5847’s tumor to 20% in P6342’s tumor. For the remainder of the analysis, we used the G0/G1 phase and normal cells to identify the underlying subclonal architecture.

We calculated the chromosomal instability (**CIN**) of the tumor cells in the G0/G1 and S phases compared to the normal cells. We determined each cell’s proportion of large-scale CNV gains and losses [27–29]. We observed different CIN levels among G0/G1 and S phase cells across all samples. The average CIN fraction in normal cell populations ranged from 30% to 50%, but the extent of CIN for cells in the S phase was much more significant, with 90% across all samples. This difference was a result of the ongoing DNA replication in dividing cells. For each case, the CIN percent of G0/G1 cells was greater than normal and smaller than replicating cells (**Figure 1D**). In contrast, cells in G0/G1 demonstrate less variation and a more stable CNV pattern [1]. Therefore, our subsequent analysis of the tumor CNVs was based on cells in G0/G1.

### Comparing chromosomal arm aneuploidy events with other cancer data sets

We compared the patterns of CNV changes on chromosomal arms among different data sets. This comparison used the G01/G1 single-cell results versus published results from surveys of primary gastric cancers. For this comparison, we used the CNVs from the stomach adenocarcinoma (**STAD**) samples reported in the Cancer Genome Atlas (**Methods**). Here, we observed concordant patterns of genomic aberrations on some chromosomal arms. Between the two data sets, there were seven overlapping chromosomal arms among the top 10. The most frequent arms with amplification events included chromosomes 5p, 7(p, q), 8(p, q), and 20(p, q) (**Figure 1E**). Meanwhile, deletion imbalances in chromosomes 9p, 17p, 18q, 19p, 21(p, q), and 22q were common. Citing an example, a deletion of the 17p arm has been demonstrated to be a driver of gastric cancer. This chromosomal alteration was evident among the gastric cancers analyzed with scDNA-seq and the TCGA data [31].

### Single-cell clonal architecture of primary gastric cancers

We applied our analysis pipeline to all cancers in our cohort (**Methods**). As already noted, we used only the data from the G0/G1 cells for identifying cancer-related CNVs. We discuss the result from P5931’s primary gastric cancer to illustrate the results of subclonal structure (**Figure 2**). Approximately half of the cells (~47%) were identified as normal cells showing diploidy. Approximately 10% of cells were detected as replicating cells in the S phase. We utilized the UMAP and Louvain clustering algorithms embedded in Seurat pipeline. Cell clusters identified distinct cell populations based on CNVs. The phylogenic output was used to define clusters as subclones.

**Figure 2.**
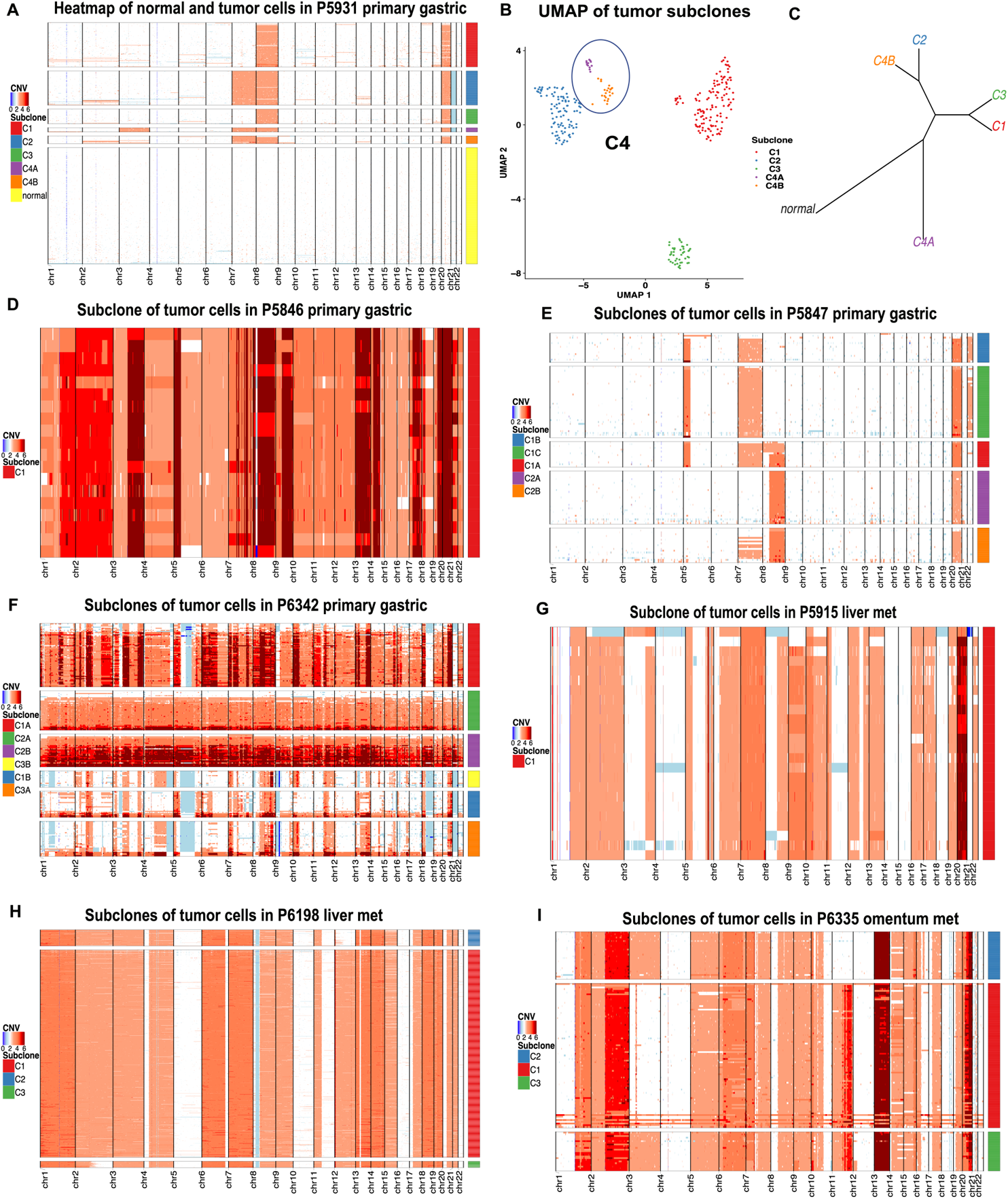
Constructed subclone architectures of scDNA-seq primary gastric and CRC liver/omentum metastasis samples. (A) Heatmap of subclone structure from G0/G1 tumor cells with detected normal cells in P5931 primary gastric. (B) UMAP plot for constructed subclone clustering of P5931 G0/G1 tumor cells. (C) Phylogenetic branches of subclones in P5931. (D-F) Heatmap plot of tumor subclone structures for three primary gastric cancer samples: P5846, P5847, P6342, respectively. (G-I) Heatmap plot of tumor subclone structures for three CRC metastasis samples: P5915 liver metastasis, P6198 liver metastasis and P6335 omentum metastasis, respectively.

P5931’s gastric cancer was dominated by significant copy number alterations affecting chromosomes 2, 3, 7, 8, and 20 (**Figure 2A**). The G0/G1 tumor cells aggregated into four major cell clusters (C1 - C4) (**Figure 2B**). Then, we applied conventional K-means clustering methods to identify minor clusters (**Methods**). For example, there were two subclusters C4A and C4B within C4.

We used a phylogenic approach to assign clusters as specific subclones. For P5931’s gastric cancer, there were six branches which represented five subclones. One branch represented the normal diploid cells. The remaining five branches were defined by the cancer CNVs. For example, Chromosome 3 amplification was the defining somatic event that separated the C4A subclone from the others (**Figure 2C**).

The three subclones (C1, C2 and C3) were defined by CNVs in Chromosomes 7, 8, and 21 (**Figure 2A, B**). The C4A and C4B subclones were dominated by chromosome 3 amplification and chromosome 21 deletion (**Figure 2C**). As previously reported, the subclone structure in P5931’s tumor was validated with other methods [30]. Thus, we determined that our approach to defining subclones was reproducible.

Next, we applied the scDNA-seq analysis pipeline to the remaining three primary gastric tumors. The results showed the subclonal diversity across these stomach cancers. For example, there was only one subclone in P5846’s gastric cancer. For this tumor, nearly all chromosomal regions had a significant number of large CNVs (**Figure 2D**). These features were indicators of general aneuploidy. This cancer had a small number of cells. Thus, the single subclone may reflect a geographically limited distribution of the tumor biopsy.

For P5847’s gastric cancer, four chromosomes demonstrated significant amplification changes including the 5q arm. CNVs were identified in chromosomes 7, 8, 20, and 22 (**Figure 2E**). Based on the cell clustering and phylogenic branch analysis, there were two major cell clusters (C1 and C2). The phylogenic analysis identified a total of five subclones. Two were dominated by changes in chromosomes 8 and 22. The remainder were defined by amplifications in Chromosome 5 and 7 (**Figure 2E and Additional File 1: Figure S5 A**). The presence of smaller, minor subclones was defined by changes in Chromosomes 8 and 22. In addition, a subset of cells that had also acquired Chromosome 7 amplifications. This distinction represented a potential intermediate step between the C2B subclone and the C1A subclone.

P6342’s gastric cancer had the most complex subclonal structure. All chromosomes showed evidence of large CNV segments. A proportion of the cells showing extensive aneuploidy over the entire genome and other cell populations had restricted chromosomal amplifications or deletions (**Figure 2F**). These CNV features define three major clusters. The phylogenic analysis identified a total of six subclones (**Additional File 1: Figure S5B**). Three subclones (C1A, C2A and C2B) showed widespread aneuploidy changes. Three other subclones (C3B, C1B and C3A) showed more discrete CNVs involving smaller segments of the genome or chromosome-arm specific changes.

### Single-cell clonal architecture of metastatic colorectal cancers

Among the colorectal cancers, we performed scDNA-seq on two liver metastasis (P5915 and P6198) and one omentum metastasis (P6335). We analyzed their single-cell copy number profiles and resolved the subclonal structure of each tumor.

For P5915’s CRC liver metastasis, there was a single subclone with 22 cells. The chromosomal-level amplifications were extensive and were evident among 12 chromosomes including 2, 6, 7, 9, 10, 12, 13, 16, 17, 19, 20, 22. Four arm-level amplification events occurred in these subclones, including 1q, 3q, 5p and 8q (**Figure 2F**). Moreover, we observed that almost 83.5% of this sample’s cells were diploid. There was a small proportion of aneuploid cells distributed in G0/G1 phase (11.3%) and S phase (5.2%) (**Additional File 2: Table S4**).

For P6198’s CRC liver metastasis, we identified three major clusters (C1, C2 and C3). The number of cells in the C2 and C3 clusters was less than C1. These clusters were assigned to three phylogenetic branches, indicating they were subclones (**Figure 2H and Additional File 1: Figure S5 C**). For the C1 subclone, amplifications cover 17 chromosomes and one chromosome arm. The smaller C2 and C3 subclone populations were distinguished by CNVs in Chromosome 12 and 2q arm, respectively (**Figure 2H**).

P6335’s CRC omentum metastasis separated into three major clusters (C1, C2 and C3) (**Figure 2I**). The three clusters were located at three distant branches by phylogenic analysis and thus assigned as distinct subclones (**Additional File 1: Figure S5 D**). Multiple whole chromosome and arm-level amplifications were present in all subclones (**Figure 2I and Additional File 1: Figure S5 D**). Amplifications on chromosomes 3 and 11 dominated the three subclones.

### Single cell DNA-seq analysis of matched colorectal cancers and distant metastasis

For two patients (P6461 and P6593), we had a subset of CRCs with matched paired primary colorectal and liver metastases. We conducted the scDNA-seq on these paired samples and studied the differences between the primary tumor originating in the colon versus the metastasis.

#### P6461 matched primary and liver metastasis

For P6461’s CRC, we analyzed a paired primary and liver metastasis. We identified 179 diploid and 245 aneuploid cells phase from the primary colorectal tumor sample. However, the metastatic tumor had only five diploid cells and six aneuploid cells **(Additional File 2: Table S4)**. The low number of cells in the liver metastasis resulted from prior chemotherapy.

This CRC had a diverse set of subclones, demonstrating a broad cellular divergence. Based on the CNV segments, we observed multiple major clusters (C1-C6) of the primary tumor and one individual metastatic cluster (**Figure 3A**). There were four major branches. For the primary colon cancer, the phylogenic analysis identified eight subclones. The metastatic tumor had only one subclone (**Figure 3B**). One branch contained the C2 subclone and the normal diploid cells. The C2 subclone had an amplification of the chromosome 19q arm. The remaining seven subclones had a broader set of chromosome alterations. In total, these CNVs covered large segments involving chromosome 6, 7, 8, 10, 12, 13, 14, 17, 18, 19, 20 and 21 (**Figure 3C**).

**Figure 3.**
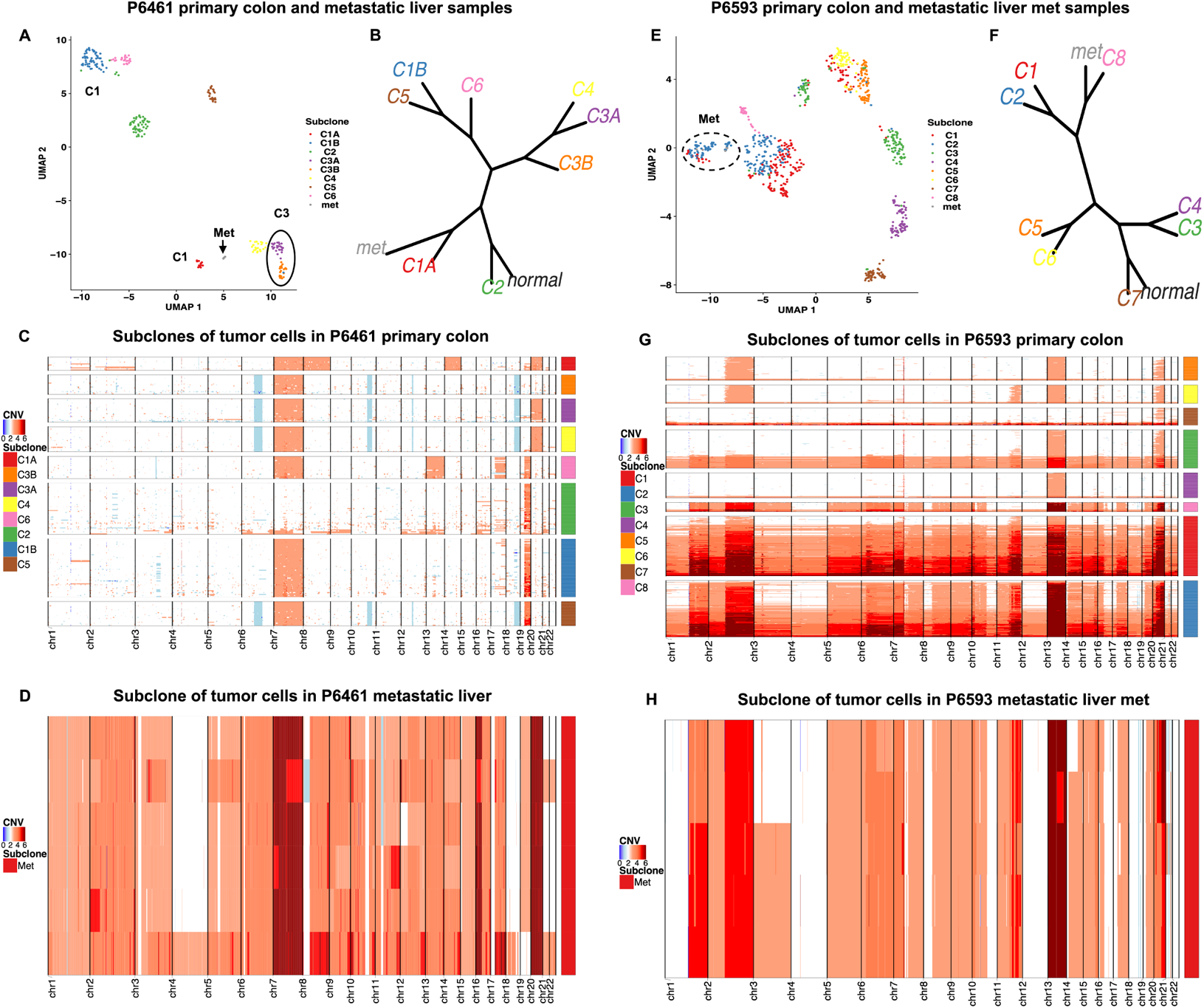
Comparing clonal architectures between paired primary colorectal cancer and metastatic liver cancer samples. (A) UMAP plot multiple subclones from primary colon with one metastasis subclones for P6461 patient. (B) Phylogenetic branches among primary colon subclones with the metastatic liver subclone in P6461 patient. (C) Heatmap plot G0/G1 tumor subclonal structures in P6461 primary colon sample. (D) Heatmap plot G0/G1 tumor subclonal structures in P6461 metastatic liver metastasis sample. (E-H) The same illustration with (A-D) for the P6593 tumor.

For the primary CRC, a set of subclones (C3A, C3B, C4 and C5) had a similar complement of amplifications but also had large segmental deletions. They all had amplifications on chromosomes 7. However, C3A and C4 had amplifications of chromosome 20 that were not observed among the others. These subclones were notable for the presence of large segment deletions involving chromosomes 6, 10, 12 and 18q. The 18q deletion is notable since that has been observed frequently and is associated with loss of *SMAD4*.

For the primary CRC, one branch contained the C1B, C5 and C6 subclones – they were defined by large segmental amplifications involving chromosomes 7 and 19q. The C6 subclone had Chromosome 13 amplification. The C5 subclone diverged with large segmental deletions as already described. The C6 subclone had exclusive large amplifications affecting chromosomes 13 and 17q but lacked the presence of the large deletions observed among some other subclones.

Finally, the metastatic subclone and C1A were on identical branch **(Figure 3B)**. Amplifications of chromosomes 8 and 14 defined the C1A subclone. The liver metastasis had general aneuploidy features across all chromosomes **(Figure 3D)**.

#### P6593 matched primary and liver metastasis

For P6593’s CRC, the primary tumor had 802 diploid cells and 759 G0/G1 aneuploid cells. The metastasis had 208 diploid cells and 5 G0/G1 cells with aneuploidy (**Additional File 2: Table S4**). The copy number data from both samples were aggregated. We detected eight major clusters (C1-C8) from the primary tumor and one cluster from the matched liver metastasis (**Figure 3E**). The eight major clusters were placed on four major branches (**Figure 3F**). Each branch had individual subclones, represented as sub-branches.

For P6593’s primary tumor and metastasis, two branches were located closer to the normal cells (**Figure 3F**). The C7 subclone originating from the primary tumor, occurred on a branch that was associated with the normal cells. CNV changes in chromosome 20 defined this subclone. The other major branches were separate from the normal cells and defined subclones with more extensive genomic instability changes. Two branches defined four subclones (C3, C4, C5 and C6) which also originated from the primary tumor. These clones were defined by changes in chromosomes 13 and chromosomal arm changes in 2p and 21q.

There were two sub-branches from C1 and C2, representing two additional subclones in the primary CRC (**Figure 3F**). These clusters were notable for having the highest number of CNVs found across all chromosomes. The remaining branch had two sub-branches defining the metastasis and a primary CRC cluster (C8). The subclones in this sub-branch showed widespread aneuploidy changes affecting all chromosomes (**Figure 3G, H**). They had the common amplification features on chromosomes 2q and 13 that were observed in all the other primary subclones (**Figure 3G, H**). This integrated analysis points to the metastasis having originated from primary tumor cells with a higher degree of aneuploidy.

### Single cell gene expression and inferred CNV clonal assignment

We performed scRNA-seq on the same set of gastric and colorectal cancers that were used for the scDNA-seq analysis (**Table 1, Additional File 2: Table S1**). The samples included patient-matched tumor, normal tissue, and metastatic tissue. Notably, the RNA-seq data sets originated from the same cell suspensions used for the scDNA-seq.

The sequencing metrics are summarized in **Additional File 2: Table S2**. Most single cells had high-quality data except for P5847’s normal tissue sample. We applied QC filtering (**Methods, Additional File 2: Table S4 and S5**). Following these procedures, we merged the remaining high-quality data, performed dimension reduction, and made cell cycle assignment and clustering (**Methods**). Finally, the Seurat clusters of the merged objects per patient were visualized by UMAP plotting, as well as the cell type biomarkers expression patterns across clusters were shown by heatmaps (**Additional File 1: Figure S6 A-C**).

We identified the clusters with the epithelial cell type by calculating the module score of relative canonical markers genes (**Methods, Additional File 2: Table S6**). Then, we extracted the epithelial cells for subsequent analysis. This processing included unsupervised cell clustering with Seurat. The results were visualized with UMAP plots (**Additional File 1: Figure S6 D, E**). The epithelial cells expressed specific epithelial markers (**Additional File 1: Figure S6 F**).

We selected the epithelial cells from G0/G1 phase by cell cycle assignment scores. We used the inferCNV package to estimate the inferred large-scale CNVs. From the scRNA-seq data, the inferred CNVs were estimated based on the relative gene expression in comparison with reference cells (**Methods**). We determined which epithelial cells had somatic CNVs, with this feature being an indicator of cancer cells. Also, this step of the analysis utilized a correlation algorithm based on an assumption of gene dosage being associated with CNVs (**Methods**). The CNV matching process was the basis for the subclonal assignment among the cancer epithelial cells identified by scRNA-seq. Also, we identified the other cell types, including fibroblasts, endothelial, macrophages, dendritic, B plasma and T cells (**Figure 4A**). The other cell types provided control for inferred CNVs.

**Figure 4.**
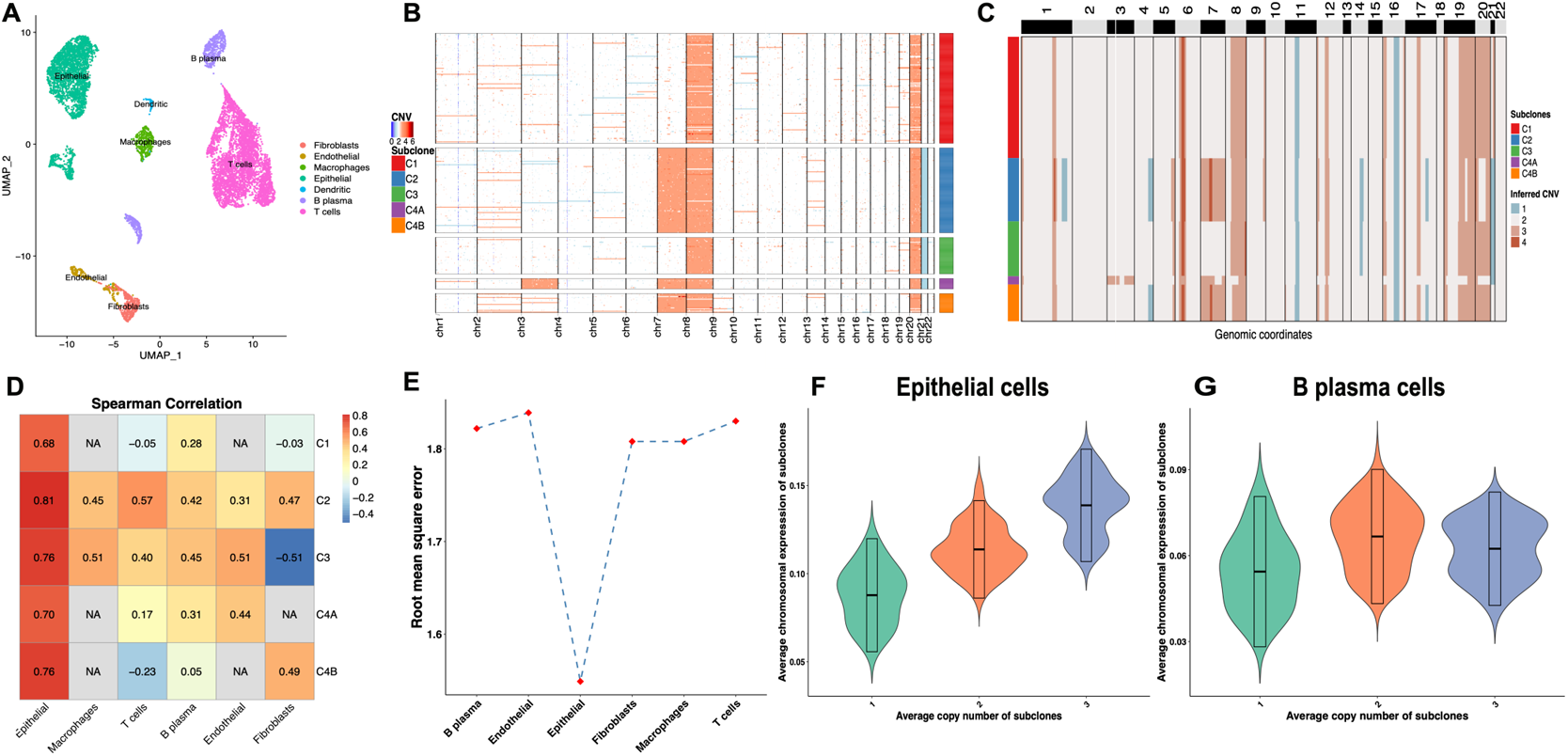
Single cell RNA-seq properties and copy number inference. (A) UMAP plot the cell type clustering and annotation of P5931 single-cell RNA-seq data. (B) Heatmap plot of the subclone structure of P5931 scDNA-seq tumor G0/G1 cells. (C) Heatmap plot inferred copy number change patterns of assigned subclones from P5931 epithelial cells scRNA-seq data. (D) The heatmap shows Spearman correlation of the CNVs from the scDNA-seq subclones with inferred CNVs from respective scRNA-seq subclones across different cell types. (E) The root mean square error (**RMSE)** calculated by the subclonal inferred CNV profile of scRNA-seq with the ground truth scDNA-seq subclonal CNVs. (F-G) Comparing the violin plots of epithelial cells B plasma cell. Each plot represents the trend between average chromosomal gene expression of scRNA-seq subclones with the corresponding average chromosomal CNVs of scDNA-seq subclones.

We use P5931’s gastric cancer to illustrate the multi-omics integration and its results. As shown in **Figures 4B and C**, the inferred cancer epithelial CNVs from scRNA-seq were generally consistent with the CNVs per the scDNA-seq analysis. As a control, the copy number integration procedure was applied to these other cell types. The heatmap of Spearman correlation across cell types showed that epithelial cell’s inferred CNV values. Among the various cell types, epithelial cells had the highest correlation with their matched scDNA-seq somatic CNV calls (**Figure 4D**). As another test, we determined the smallest root mean square error (**RMSE**) based on the inferred CNV profile for each profile. We used scDNA-seq CNVs as the ground truth. The epithelial cell CNVs per scRNA-seq had the lowest squared errors compared with the other cell types (**Figure 4E**). We examined the dosage effect between genome and transcriptome on matched subclones. The average of chromosomal scaled gene expression of the subclones showed a positive correlation with the average CNV of scDNA-seq subclones. We calculated the average gene expression on each chromosome in a scRNA-seq subclone versus the average CNV on the same chromosome of the corresponding scDNA-seq subclone. We used the Chi-squared test for spearman correlation. The epithelial cell’s p-values were 2.66e-3 (*AvgExp on CN = 1 and CN = 2*) and 2.14e-7 (*AvgExp on CN = 2 and CN = 3*) (**Figure 4F**). In contrast, the B cells did not show any trends of average expression between average CNV=2 and CNV=3, as well as other cell types. Moreover, for these other cell types, the Chi-squared correlation test had no significant difference between average expression when form copy number 2 to 3 (**Figure 4G**). The same trend of a lack of correlation was noted across all non-epithelial cells.

Finally, we demonstrated that the correlation algorithm could accurately give the clonal assignment of the epithelial cells (per the scRNA-seq data). This step involved a permutation test. The observed RMSE in the clonal assignment was significantly smaller than the computed RMSEs under random permutation of clone assignments (**Additional File 1: Figure S6 G**).

### Pathway analysis of subclonal populations in gastric cancer

We identified the biological pathway features from those tumors with multiple subclones. This involved using a GSVA pathway analysis on the epithelial cells identified from the scRNA-seq data. We made a series of pairwise comparisons among the different subclones and the epithelial normal cells when available. Differences in the activities of canonical pathways among the various epithelial cellular subpopulations, such as normal versus tumor, primary versus metastasis tissues, were determined. We detail a list of some relevant examples with high statistical significance (p < 1.00E-07) is shown in **Table 2** and provide some heatmaps of specific tumors (**Figure 5**). A full compilation of the most significant pathway features of subclones are provided in **Additional file 2: Table S8**.

**Figure 5.**
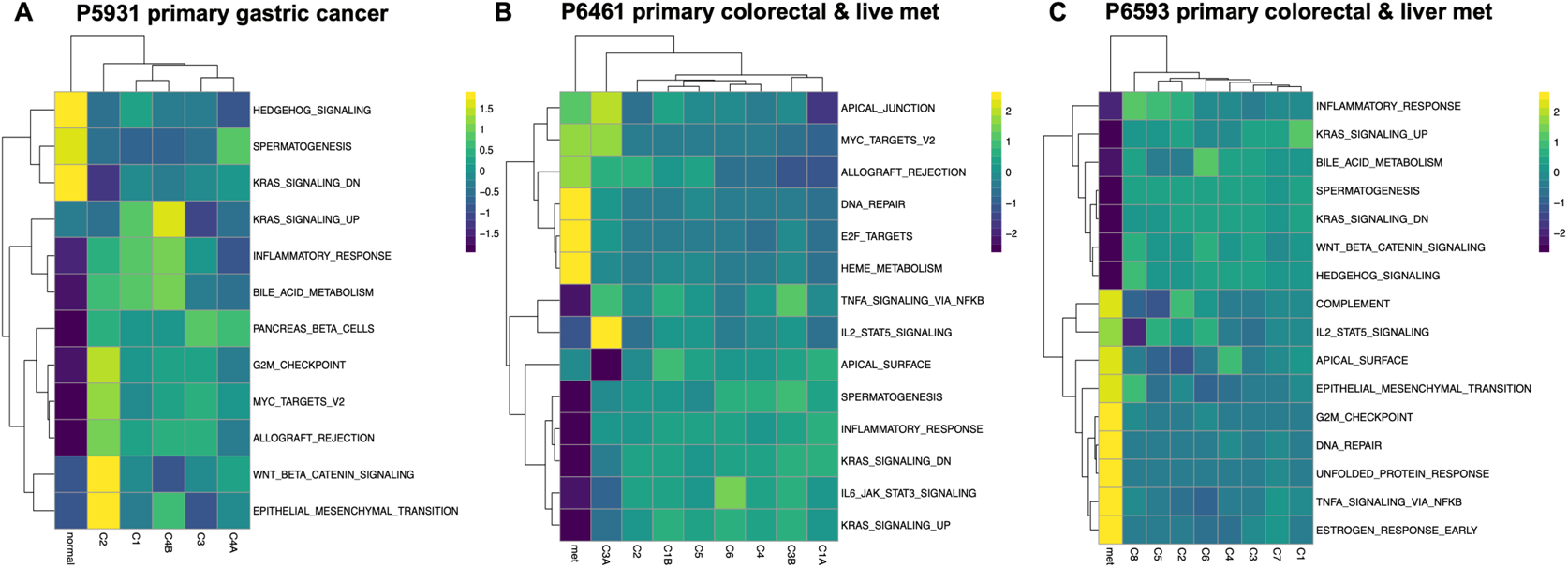
Heatmap depicting average scaled GSVA enrichment score of selected Hallmark pathways for scRNA-seq subclones. (A) P5931 primary gastric cancer. (B) Matched primary colorectal sample and metastatic liver metastasis sample from P6461 patient. (C) Matched primary colorectal sample and metastatic liver metastasis sample from P6593 patient. Significance was based on an ANOVA Tukey HSD analysis (*p-value < 0.05).

**Table 2.** Examples of significant pathway differences among subclones.

| Cancer type | Patient ID | Subclone | Pathway |
| --- | --- | --- | --- |
| Gastric | P5847 | C1A | bile acid metabolism |
|  |  | C1C | TNF-alpha signaling via NFkB |
|  |  |  | Notch signaling |
|  | P5931 | C2 | epithelial mesenchymal transition |
|  |  |  | G2M checkpoint |
|  |  |  | Wnt signaling |
|  |  | C4b | KRAS signaling |
|  | P6342 | C1A | heme metabolism |
|  |  |  | KRAS signaling |
|  |  |  | TNF-alpha signaling via NFkB |
|  |  | C3A | epithelial mesenchymal transition |
|  |  |  | E2F transcription pathway |
|  |  |  | G2M checkpoint |
|  |  | C3B | angiogenesis |
| Colorectal | P5915 | C1 | TP53 pathway |
|  | P6198 | C1 | TNF-alpha signaling via NFkB |
|  | P6335 | C1 | AKT signaling |
|  |  |  | Wnt signaling |
|  |  |  | Myc signaling |
|  | P6461 | metastasis | E2F transcription pathway |
|  |  |  | DNA repair |
|  |  | C3A | IL 5 Stat signaling |
|  | P6593 | metastasis | DNA repair |
|  |  |  | epithelial mesenchymal transition |
|  |  |  | G2M checkpoint |
|  |  |  | hedgehog signaling |
|  |  |  | KRAS signaling |

Among the primary gastric cancers, the different subclones had diverse pathway activity (**Additional File 2: Table S8**). As an example, we describe some of the notable subclonal features from P5931’s gastric cancer (**Figure 5A**). Two of the subclones had distinct pathway features that denoted them from the others. The C2 subclone had distinct cancer pathways with a high level of upregulation including the Wnt signaling pathway and the epithelial mesenchymal transition. The C2 subclones had a deletion of chromosome 21 as one of its defining genomic DNA features. The C4B subclone had elevated KRAS pathway activity that was significantly higher than the other subclones and normal epithelial cells.

For the P5847 primary gastric cancer, the majority significant pathways enriched in C1 subclone included Notch signaling and TNFA signaling via NFKB (**Table 2**). The C1A subclone had pathway activity in bile acid metabolism.

For the P6342 gastric tumor, the C1A subclone had higher KRAS activity and TNFA signaling via NFKB activity. The C1A subclone had a high degree of aneuploidy affecting all chromosomes (**Figure 2F**). The C3A subclone had higher activity in the epithelial mesenchymal transition, E2F transcription and G2M checkpoint pathways. The C3B subclone had higher activity in the angiogenesis pathway. The C3A and C3B subclones did not have the degree of aneuploidy noted in C1A.

### Pathway analysis of subclonal populations in metastatic colorectal cancer

Next, we evaluated the subclones from the matched primary colorectal cells with live metastasis (P6461 and P6693). There were several significant pathways with higher GSVA scores present in the liver metastasis subclone (**Figure 5**). For P6461’s metastasis, three significant pathways were involved in the liver metastasis cells comparing with other primary colorectal subclones (**Figure 5B**). They included DNA repair, E2F-transcription factor signaling and heme metabolism. There was only one significant pathways in subclone C3A, which showed IL2-STAT5 signaling activity.

For P6593’s subclones, the liver metastasis had significant upregulation of multiple pathways compared to the other subclones making up the primary CRC in the colon (**Figure 5C**). Some of the notable pathways included increase activity in the epithelial mesenchymal transition, G2M checkpoint, E2F-transcription factor signaling, MTOROC1 signaling, MYC signaling, and DNA repair. The differences were less pronounced among the subclones in the primary CRC. However, the C8 subclone did have elevated activity in the epithelial mesenchymal transition pathway compared to the other subclones. The C8 subclone was also on the same major branch as the metastasis. The CNV patterns of the C8 subclone showed more widespread genomic instability compared to the other primary CRC subclones. These results suggest that the C8 may have had some biological link to the metastasis with biological properties that enable it to spread to the liver.

For the P6335 omentum metastasis, the downregulated UV response, apical surface, and inflammatory response activities were involved in normal cells. The tumor subclone C1 had several pathways enrichment, including Wnt beta catenin signaling, p13k AKT Mtor signaling as well as MYC targets signals. Reviewing the other CRCs, for P5915’s tumor, there was only a single major subclone. For the P6198 CRC, the C1 subclone had TNFA signaling via NFKB.

## DISCUSSION

Single-cell multi-omics provides a wealth of information on the cellular features of cancer. Cancers have multiple subclones that have distinct genomic and cellular properties. An important topic of cancer research is determining the phenotypic properties of these subclones and their biological features. This cellular variation is essential in facilitating metastasis and provides a basis for resistance to specific therapies. A variety of single-cell genomic assays provide specific cellular and genomic characteristics. However, the use of a single type of assay has limitations. For example, scDNA-seq provides genomic DNA variation such as CNVs at single-cell resolution and characterizes subclone architectures of tumors. However, scDNA-seq will not identify the type of cell or its phenotype. Single cell RNA-seq provides a wealth of gene expression information that elucidates an individual cell’s biology. However, its use in defining specific cancer subclones is limited, given the intrinsic issues of extrapolating clonal structure from gene expression patterns. For instance, the calling of CNVs from scRNA-seq remains an area that would benefit from improvements. By integrating both scDNA-seq and scRNA-seq, one defines subclonal structure based on genomic DNA variation, identifies single-cell gene expression patterns, and determines pathway activity for specific cancer cell subpopulations.

This multi-omics approach successfully integrated scDNA-seq and scRNA-seq analysis on a set of gastric and colorectal cancers from surgical biopsies. This approach identified both the genomic CNV variation, subclonal structure, and transcriptome phenotype of each cancer. Single-cell genotype-phenotype features reveal important malignant properties relevant to proliferation, metastasis, and treatment response in cancer [1, 11]. This multi-omics approach may prove useful in identifying specific biomarkers indicative of subclonal populations that influence the clinical course of patients.

To integrate these data sets, we identified the cells in G0/G1 phase and replicating cells in the S phase. Distinguishing these populations based on their position in the cell cycle improved the quality of our results. Determining how the cell cycle affects the quality of the subclonal analysis has not been thoroughly investigated [15, 32]. Our prior study of gastric cancer cell lines showed the significance of these effects [1]. This study demonstrated that the analysis of primary cancers from surgical biopsies. Analyzing tumor cells in G0/G1 phase improved the reconstruction of subclonal architecture. Thus, this study focused on epithelial cells in G0/G1 phase from both data sets.

Integrating the DNA and RNA features were based on the gene dosage effects and their consequences on expression [17]. Copy number amplification increases gene expression, while deletions can decrease expression [1, 15, 18]. To overcome the stochasticity of gene expression counts, we applied scaled gene expression to calculate the correlation with the consensus subclone CNV composition determined by scDNA-seq.

Our results provided an informative insight into these tumors. For example, primary colon tumors had greater degrees of intracellular heterogeneity than liver metastasis. Both P6461 and P6593 primary colorectal cancer samples had multiple distinct subclones (**Figure 3A, E**). Their matched metastases had only one subclone with six (P6461) and five (P6593) G0/G1 tumor cells, respectively. Interestingly, the metastases showed extensive aneuploidy. The prior chemotherapy treatment could explain the lower number of tumor cells in metastasis from the liver before resecting the tumor. Interestingly, tumor cells were present despite prior chemotherapy treatment.

## CONCLUSIONS

We used an integrative approach to leverage scDNA-seq and scRNA-seq for characterizing the genotype-phenotype characteristics of subclones within a given tumor. We identified the proportions of normal cells with diploids, G0/G1 cells, replicating cells in each tumor scDNA-seq sample, and detected the subclonal architecture of aneuploid cells in G0/G1 phase. Moreover, we assigned the epithelial cells of the scRNA-seq profile to the corresponding scDNA-seq subclones. This assignment process revealed the phenotypic features of specific subclones and their distinct biological characteristics.

## DECLARATIONS

### Ethics approval and consent to participate

This study was conducted in compliance with the Helsinki Declaration. The Institutional Review Board at Stanford University School of Medicine approved the study protocols (IRB-44036). All patients provided written informed consent to participate.

### Consent for publication

All patients consented for publication of the de-identified results.

### Availability of data and material

The datasets generated for this study are available in the dbGAP repositories; accession numbers phs001711 and phs001818. The analysis code used for this study can be accessed at https://github.com/XQBai/Single-cell-multi-omic-integration.

### Competing interests

The authors declare that they have no competing interests.

### Funding

This work was supported by the National Institutes of Health grants [R01HG006137-04 to XB and HPJ, R35HG011292 to B.T.L., P01HG00205ESH to BL and HPJ, R33CA247700 to HPJ]. Additional support to HPJ and AS came from the MBGI-21-109-01 from the American Cancer Society, the Gastric Cancer Foundation and the Clayville Foundation.

### Authors’ contribution

X.B., B.L. and H.P.J. designed the experiments. B.L. and A.S. conducted the single cell sequencing experiments. X.B., and B.L. developed the analysis algorithms. X.B., B.L., S.M.G., and A.S. analyzed the data. X.B. and H.P.J. wrote the manuscript. H.P.J. supervised the overall project. All authors read and approved the final manuscript.

## ABBREVIATIONS

scDNA-seq: single-cell DNA-sequencing
WGS: whole genome sequencing
CNV: copy number variation
scRNA-seq: single-cell RNA-sequencing
CRC: colorectal cancer
CIN: chromosomal instability
STAD: stomach adenocarcinoma
TCGA: the Cancer Genome Atlas
WGS: whole genome sequencing
EM: Expectation-Maximization
PC: principal component
SNN: shared nearest neighbor
MS: module score
GSVA: gene set variation analysis
QC: quality control
UMAP: uniform manifold approximation and projection
EGFR: epidermal growth factor receptor

## Acknowledgements

We thank the participating patients.

## Notes

### Competing Interest Statement

The authors have declared no competing interest.

