## Supplementary Materials for "Single cell multi-omic mapping of subclonal architecture and pathway phenotype in primary gastric and metastatic colon cancers"

### INSTITUTIONS

[1]Division of Oncology, Department of Medicine, Stanford University School of Medicine, Stanford, CA, United States

[2]Department of Electrical Engineering, Stanford University, Palo Alto, CA, United States

### CORRESPONDING AUTHOR

Hanlee P. Ji

Division of Oncology, Department of Medicine – Stanford University School of Medicine

CCSR 1115, 269 Campus Drive

Stanford, CA 94305-5151

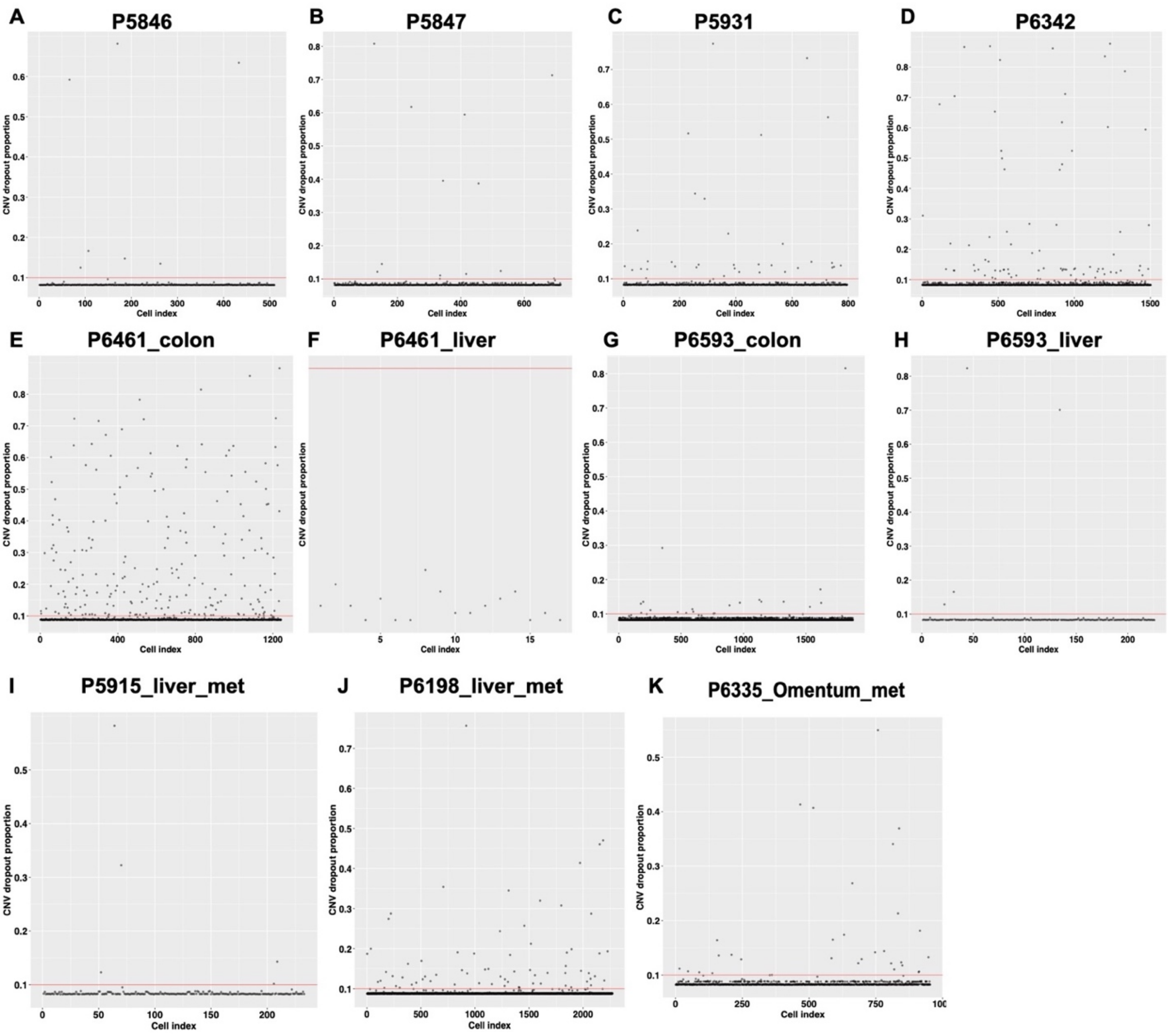


**Supplementary Figure S1**: **The proportion of CNV dropout (CN=2) for the scDNA-seq analysis of each sample.** (A-K) Scatter plot shows the dropout rate of single cells across samples. The red line is the 0.1 threshold for dropout rate. Each dot represents single cell. The dots beyond the red line were distinguished as the technical cells.

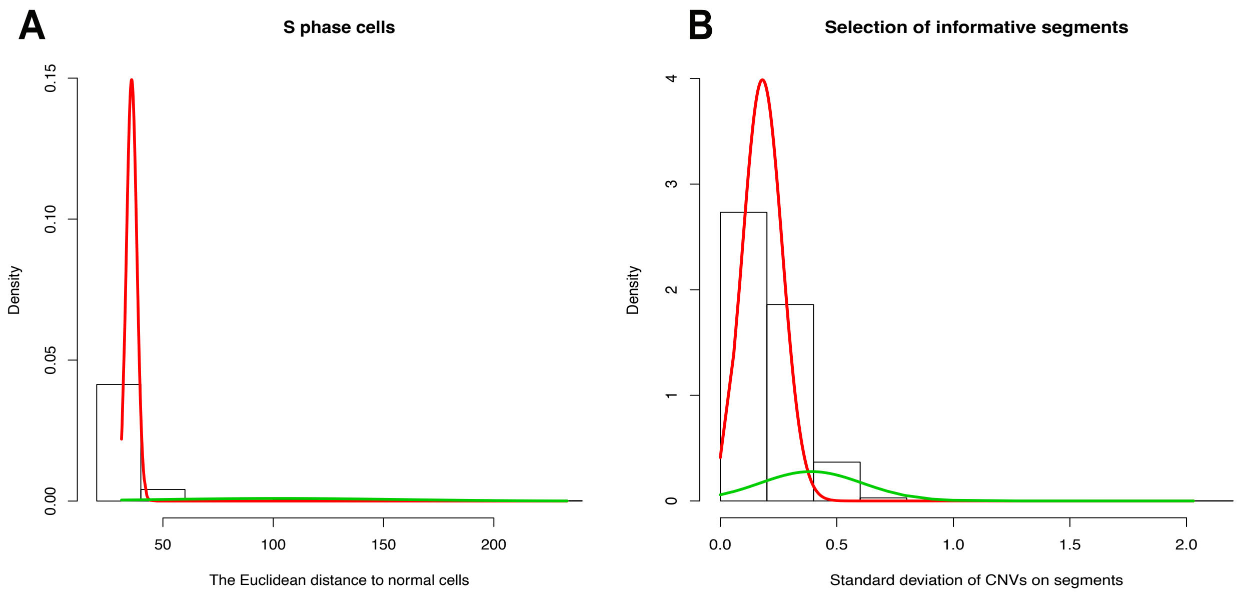


**Supplementary Figure S2: Density plot of fitted mixed normal distributions.**
(A) Modeling two mixture normal distribution for the distance from cells after filtering noise to the normal cell profile. Here, the mean of the normal distribution under red line is greatly smaller than the mean of another normal distribution with green line, which means the cells/dots located at the normal distribution with green were identified as replicating cells in S phase. (B) Fitting mixture normal distribution for the standard deviation (SD) of CNV segments across cells by EM algorithms. The CNV segments with small SD barely contribute to the subclone structures. Then the CNV segments from the normal distribution with high mean were selected as the informative scDNA-seq features to construct subclones.

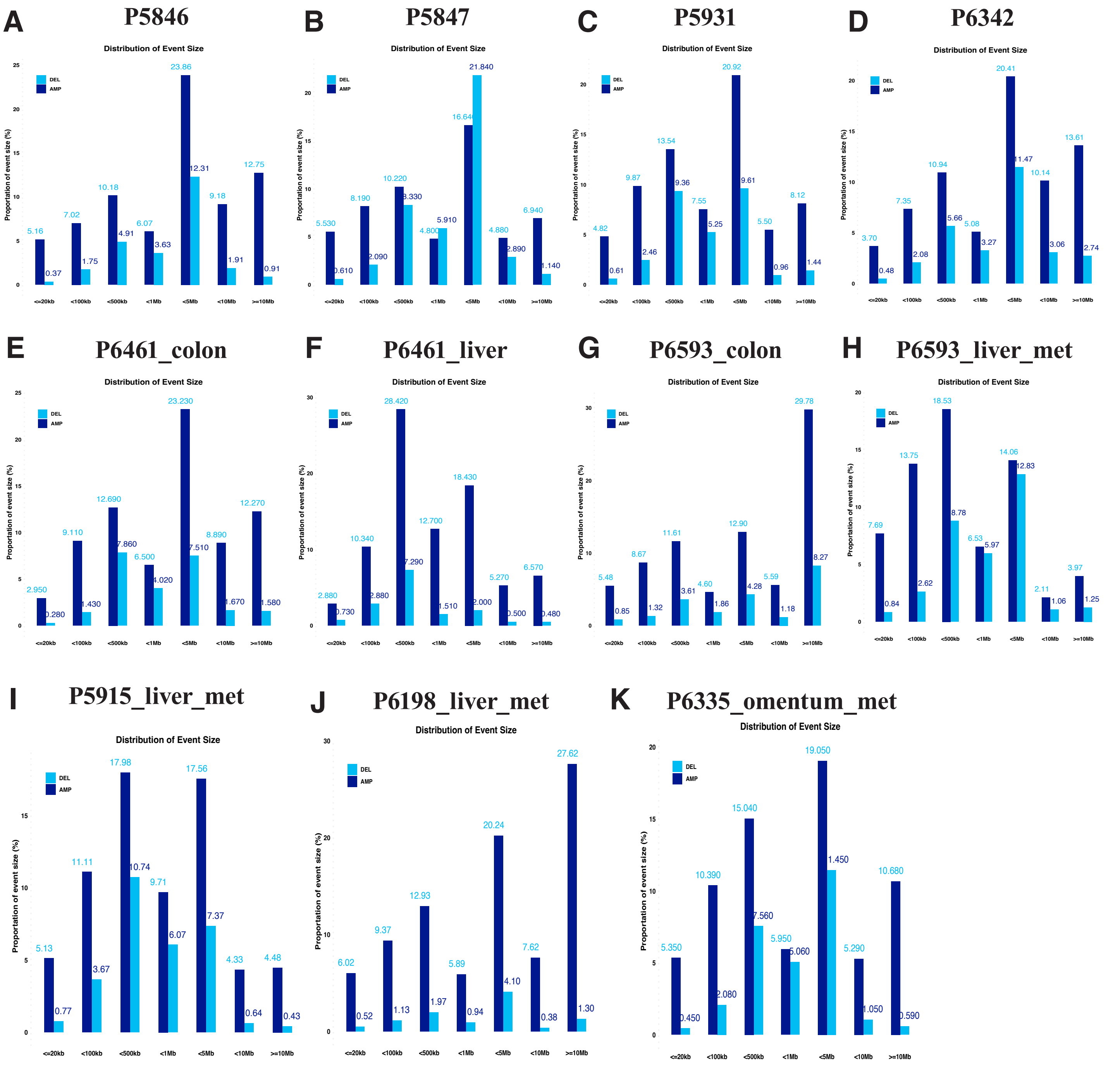


**Supplementary Figure S3: The proportion of amplification (AMP) and deletion (DEL) events across the genomic sizes.** (A-K) Across all samples, AMP (dark blue) and DEL (light blue).

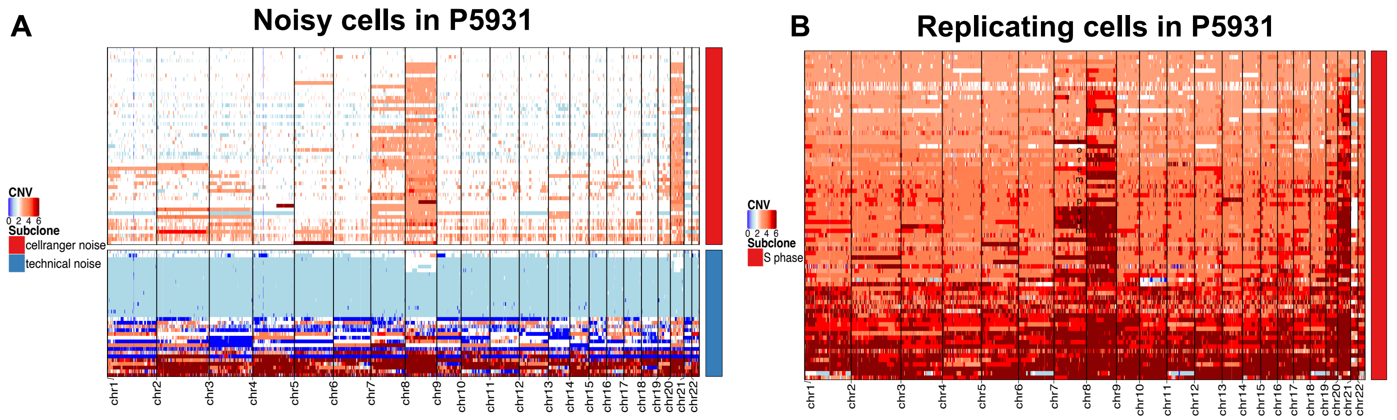


**Supplementary Figure S4: Heatmaps visualize CNV changes of noise and replicating cells in P5931.** (A) Heatmap plot the CNV profiles of noisy cells in P5931, which consists of technical noise and cellranger noise cells. (B) Heatmap plot the CNV of replicating cells in S phase of P5931.

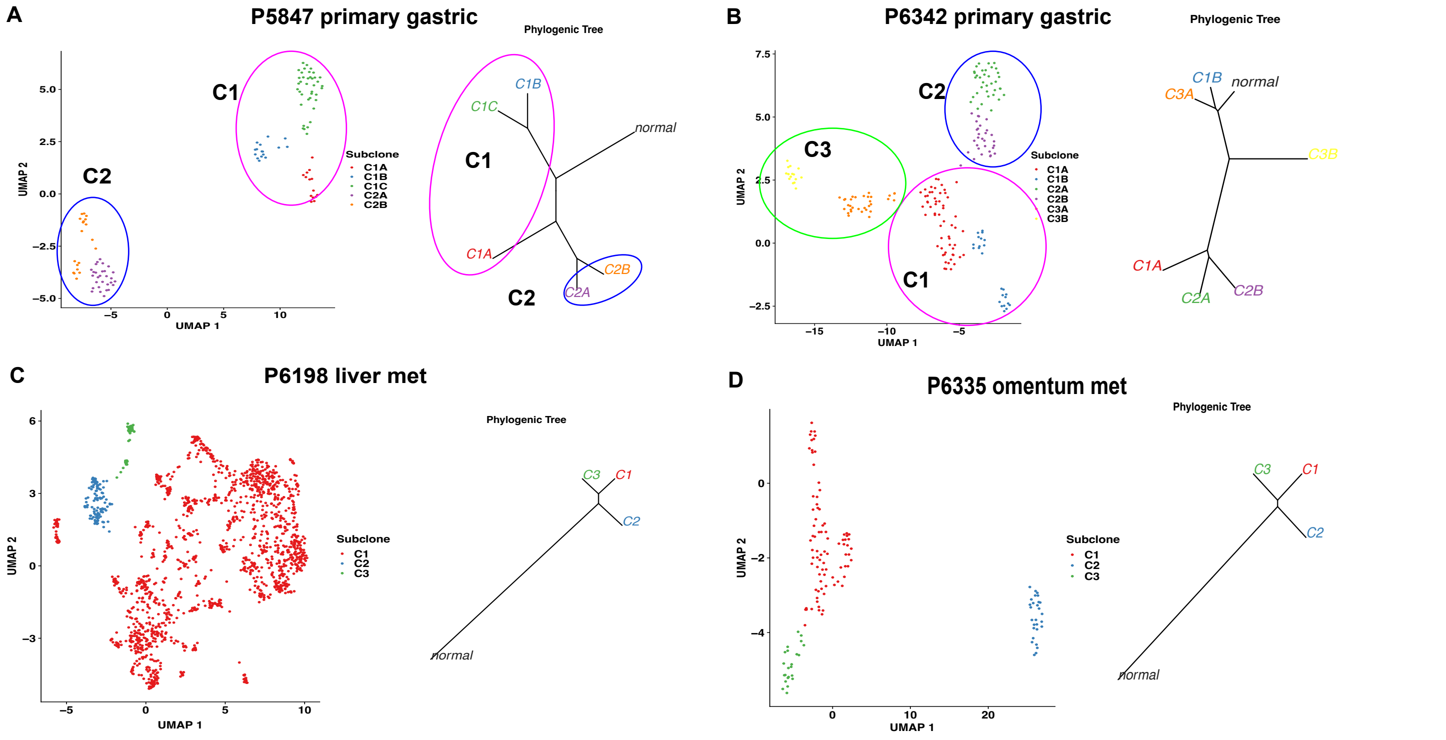


**Supplementary Figure S5: The UMAP visualize the subclones distribution of tumor G0/G1 cells and the corresponding subclonal phylogenic tree.** (A) P5847 primary gastric. (B) P6342 primary gastric. (C) P6198 CRC liver met. (D) P6335 CRC omentum met.

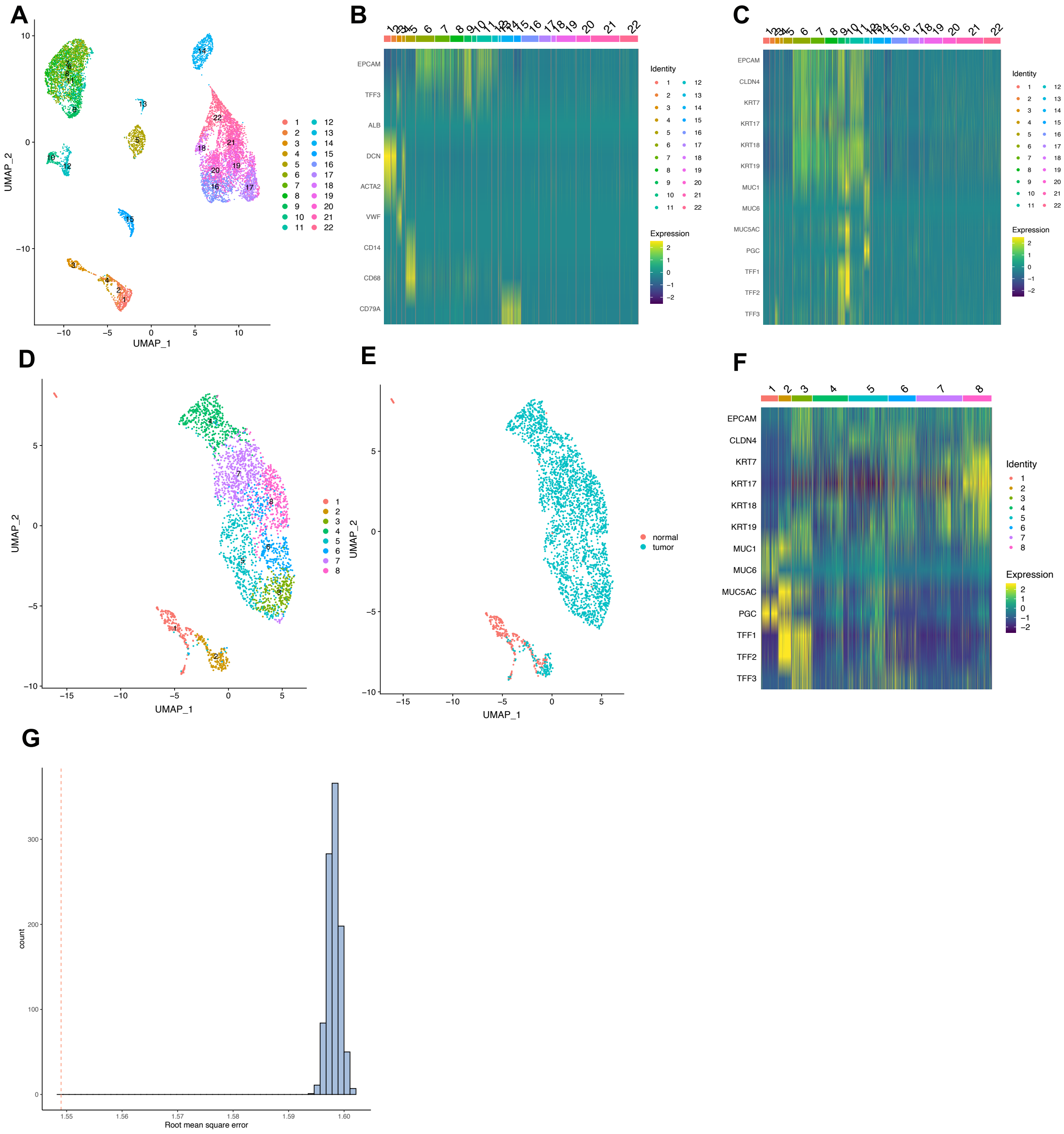


**Supplementary Figure S6: Visualization of scRNA-seq analysis for epithelial cell extraction.** (A) UMAP visualize the clusters of merged normal and tumor scRNA-seq data in P5931 by Seurat. (B) Heatmap plot the cell type biomarkers' expression patten across the clusters for cell type annotation. (C) Epithelial biomarkers expression across all clusters, in which clusters 6-12 have higher epithelial biomarkers expression, those were annotated as epithelial cells. (D) UMAP plot the re-clustering results for extracted epithelial cells of P5931. (E) UMAP shows the normal and tumor epithelial cells distribution in P5931. (F) Heatmap of epithelial biomarkers on the identified epithelial cells in P5931. (G) Permutation test for RMSE conducted for the ground truth of scDNA-seq consensus CNV of subclones with the consensus inferred CNV of assigned subclones of P5931 epithelial scRNA-seq. The red line was the RMSE calculated by scRNA-seq subclones assignment conducted as the correlation algorithm. The right distribution was the RMSE values when randomly permutated scRNA-seq subclones assignment 1000 times.

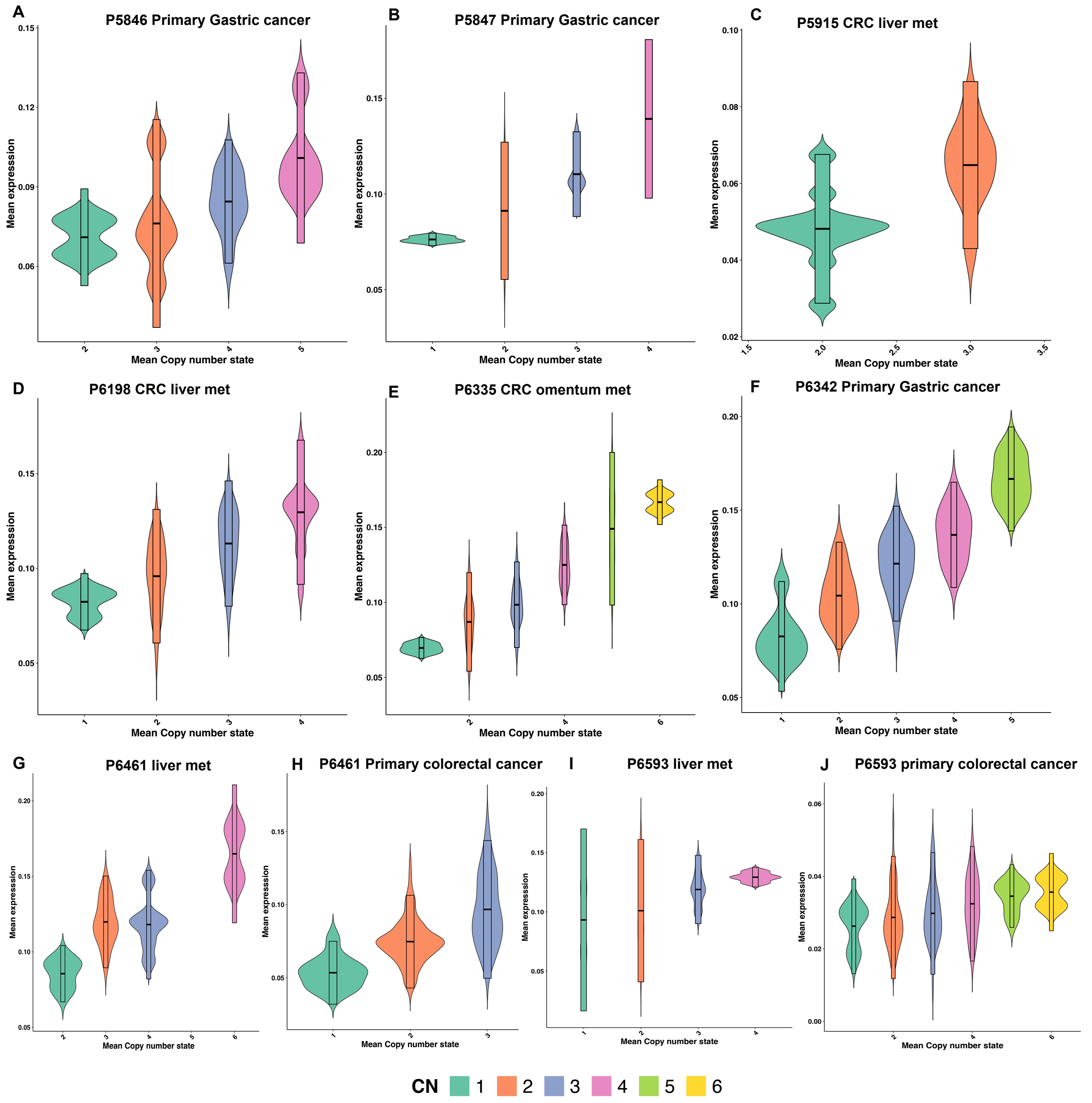


**Supplementary Figure S7: Violin plots comparing the mean gene expression distribution under the different mean copy number states of subclones.** The x-axis is the mean copy number, which was calculated as the average CNVs on chromosomes for scDNA-seq subclones. The y-axis is the mean gene expression, which was calculated as the average of chromosomal gene expression of scRNA-seq subclones. For all violin plots, the groups with different copy number variants were colored as lime green (CN=1), red orange (CN=2), polo blue (CN=3), tea rose (CN=4), yellow green (CN=5) and gold yellow (CN=6).

| Supplementary Table S1. Summary of samples | | | |
|---|---|---|---|
| **Patient ID** | **Sample ID** | **Tissue Source** | **Tissue Pathology** |
| **P5846** | P5846_18344_normal | Stomach | Normal tissue |
| | P5846_18345_tumor | Stomach | Cancer |
| **P5847** | P5847_18349_normal | Stomach | Normal tissue |
| | P5847_18350_tumor | Stomach | Cancer |
| **P5931** | P5931_800_normal | Stomach | Normal tissue |
| | P5931_801_tumor | Stomach | Cancer |
| **P6342** | P6342_19538B_normal | Stomach | Normal tissue |
| | P6342_19539B_tumor | Stomach | Cancer |
| | P6342_19539G_tumor | Stomach | Cancer |
| **P5915** | P5915_18363B_normal | Liver | Normal tissue |
| | P5915_18364B_tumor | Liver | Cancer |
| **P6198** | P6198_18502_normal | Liver | Normal tissue |
| | P6198_18503_tumor | Liver | Cancer |
| | P6198_normal | Liver | Normal tissue |
| | P6198_tumor | Liver | Cancer |
| **P6335** | P6335_19516B_normal | Omentum | Normal |
| | P6335_19517B_tumor | Omentum | Cancer |
| | P6335_tumor | Omentum | Cancer |
| **P6461** | P6461_19617_met | Liver | Cancer |
| | P6461_tumor_liver | Liver | Cancer |
| | P6461_normal_liver | Liver | Normal tissue |
| **P6461** | P6461_19615_tumor | Colon | Cancer |
| | P6461_normal_colon | Colon | Normal tissue |
| | P6461_tumor_colon | Colon | Cancer |
| **P6593** | P6593_21330B_met | Liver | Cancer |
| | P6593_liver_met | Liver | Cancer |
| **P6593** | P6593_21332B_tumor | Colon | Cancer |
| | P6593_colon_tumor | Colon | Cancer |

**Supplementary Table S2. Sequencing metrics of single-cell DNA-seq data**

| Sample | Num cells | Total reads | Fraction of mappablity | Fraction of noisy cells | Median estimated CNV resolution |
|---|---|---|---|---|---|
| P5846_18345_tumor | 510 | 389999648 | 0.9036089 | 0.0372549 | 2.0044434 |
| P5846_18344_normal | 251 | 291288144 | 0.9034082 | 0.1195219 | 1.768457 |
| P5847_18350_tumor | 715 | 476056120 | 0.9035183 | 0.1636364 | 2.0237305 |
| P5847_18349_normal | 538 | 376462156 | 0.9035247 | 0.1524164 | 2.998584 |
| P5931_801_tumor | 796 | 897164858 | 0.9035183 | 0.3781407 | 1.3674805 |
| P5931_800_normal | 41 | 328203478 | 0.9036024 | 0.097561 | 0.5182129 |
| P6342_19539G_tumor | 1502 | 3.703E+09 | 0.9040169 | 0.1298269 | 1.9284668 |
| P6461_normal_colon | 1346 | 1.034E+09 | 0.903978 | 0.1002972 | 2.225 |
| P6461_tumor_colon | 1242 | 1.381E+09 | 0.9039975 | 0.1972625 | 2.3656006 |
| P6461_normal_liver | 726 | 521980630 | 0.9039133 | 0.0330579 | 1.8940918 |
| P6461_tumor_liver | 17 | 175879246 | 0.9039521 | 0.3529412 | 0.5073242 |
| P6593_colon_tumor | 1878 | 2.024E+09 | 0.9039003 | 0.4041534 | 6.7262207 |
| P6593_liver_met | 226 | 207296622 | 0.9039198 | 0.0221239 | 1.4668701 |
| P5915_18364B_tumor | 233 | 407442904 | 0.9035765 | 0.0901288 | 0.8919922 |
| P5915_18363B_normal | 424 | 1.031E+09 | 0.9036153 | 0.5660377 | 0.7861572 |
| P6198_tumor | 2271 | 1.536E+09 | 0.9038809 | 0.0691325 | 1.5680664 |
| P6198_normal | 352 | 255524618 | 0.9039909 | 0.0113636 | 1.6979492 |
| P6335_tumor | 963 | 688093614 | 0.9038356 | 0.0766002 | 1.6317871 |

Note: The fraction of noisy cells for tumor samples were calculated by our methods. For normal samples, the fraction of noisy cells only represent the cellranger-cnv identified noise.

| Supplementary Table S3. Sequencing metrics of single-cell RNA-seq data | | | | | | |
|---|---|---|---|---|---|---|
| **Sample** | **Num cells** | **Mean reads per cell** | **Median genes per cell** | **Number reads** | **Genes detected** | **Median UMI per cell** |
| P5846_18344_normal | 1541 | 41497 | 642 | 63947206 | 18728 | 1896 |
| P5846_18345_tumor | 3476 | 19065 | 516 | 66270205 | 19373 | 1046 |
| P5847_18349_normal | 984 | 44 | 7 | 43461 | 3285 | 8 |
| P5847_18350_tumor | 7387 | 16184 | 505 | 119558075 | 19421 | 1067 |
| P5931_800_normal_1 | 926 | 100478 | 547 | 93042673 | 17038 | 1356 |
| P5931_800_normal_2 | 755 | 182586 | 650 | 137853083 | 17119 | 1694 |
| P5931_801_tumor_1 | 5861 | 19129 | 657 | 112119187 | 21444 | 2230 |
| P5931_801_tumor_2 | 5356 | 19642 | 640 | 105205866 | 21309 | 2133 |
| P6342_19538B_normal | 1065 | 29086 | 537 | 30977146 | 16826 | 1504 |
| P6342_19539B_tumor | 2976 | 14893 | 582 | 44322838 | 19898 | 1683 |
| P6461_19615_tumor | 3475 | 35062 | 374 | 121841924 | 18454 | 1030 |
| P6461_19617_met | 258 | 77484 | 45 | 19990956 | 13098 | 498 |
| P6593_21330B_met | 696 | 39258 | 690 | 27324057 | 15056 | 1519 |
| P6593_21332B_tumor | 3939 | 41253 | 132 | 162497508 | 16488 | 232 |
| P5915_18363B_normal_liver | 2349 | 39798 | 1076 | 93486060 | 19950 | 3092 |
| P5915_18364B_tumor_liver | 3640 | 24820 | 364 | 90347245 | 19853 | 1213 |
| P6198_18502_normal_liver | 5122 | 47063 | 443 | 241058226 | 18264 | 867 |
| P6198_18503_tumor_liver | 7190 | 51399 | 2001 | 369565082 | 22593 | 6952 |
| P6335_19516B_normal_omentum | 2212 | 6725 | 564 | 14877554 | 18041 | 1101 |
| P6335_19517B_tumor_omentum | 3184 | 35488 | 604 | 112994457 | 20895 | 1448 |

**Supplementary Table S4. The number and fraction of various cell types in single-cell DNA-seq samples after filtering noisy cells**

| Sample | #Normal cells | #G0/G1 phase cells | #S phase cells | #Total cells | Normal fraction | G0/G1 fraction | S phase fraction |
|---|---|---|---|---|---|---|---|
| P5846_tumor | 411 | 19 | 60 | 490 | 0.83877551 | 0.03877551 | 0.12244898 |
| P5847_tumor | 536 | 117 | 30 | 683 | 0.78477306 | 0.171303075 | 0.043923865 |
| P5931_tumor | 334 | 301 | 74 | 709 | 0.471086037 | 0.424541608 | 0.104372355 |
| P6342_tumor | 843 | 195 | 268 | 1306 | 0.645482389 | 0.149310873 | 0.205206738 |
| P6461_tumor_colon | 179 | 245 | 477 | 901 | 0.198668147 | 0.271920089 | 0.529411765 |
| P6461_tumor_liver | 5 | 6 | 5 | 16 | 0.3125 | 0.375 | 0.3125 |
| P6593_colon_tumor | 802 | 759 | 211 | 1772 | 0.452595937 | 0.428329571 | 0.119074492 |
| P6593_liver_met | 208 | 5 | 4 | 217 | 0.958525346 | 0.023041475 | 0.01843318 |
| P5915_tumor_liver | 177 | 24 | 11 | 212 | 0.83490566 | 0.113207547 | 0.051886792 |
| P6198_tumor_liver | 109 | 1613 | 392 | 2114 | 0.051561022 | 0.763008515 | 0.185430464 |
| P6335_tumor_omentum | 686 | 136 | 58 | 880 | 0.779545455 | 0.154545455 | 0.065909091 |

**Supplementary Table S5. The parameters set in single-cell RNA-seq data filtering procedure**

| | Single Cell RNA | | Standard/Adj filtering | |
|---|---|---|---|---|
| **Sample** | **Num cells** | **Median genes** | **Num cells** | **Num Clusters** |
| P5846_18344_normal | 1541 | 642 | 1265 | 13 |
| P5846_18345_tumor | 3476 | 516 | 3315 | 18 |
| P5847_18349_normal | 984 | 7 | 28 | 8 |
| P5847_18350_tumor | 7387 | 505 | 7183 | 20 |
| P5931_800_normal_1 | 926 | 547 | 749 | 9 |
| P5931_800_normal_2 | 755 | 650 | 593 | 9 |
| P5931_801_tumor_1 | 5861 | 657 | 4101 | 15 |
| P5931_801_tumor_2 | 5356 | 640 | 3913 | 15 |
| P6342_19538B_normal | 1065 | 537 | 938 | 11 |
| P6342_19539B_tumor | 2976 | 582 | 2853 | 14 |
| P6461_19615_tumor | 3475 | 374 | 2236 | 9 |
| P6461_19617_met | 258 | 45 | 117 | 3 |
| P6593_21330B_met | 696 | 690 | 694 | 6 |
| P6593_21332B_tumor | 3939 | 132 | 3934 | 18 |
| P5915_18363B_normal_liver | 2349 | 1076 | 2186 | 18 |
| P5915_18364B_tumor_liver | 3640 | 364 | 2399 | 12 |
| P6198_18502_normal_liver | 5122 | 443 | 4914 | 33 |
| P6198_18503_tumor_liver | 7190 | 2001 | 6674 | 19 |
| P6335_19516B_normal_omentum | 2212 | 564 | 2140 | 13 |
| P6335_19516B_normal_omentum | 3184 | 604 | 2833 | 17 |

1. Few cells, low genes/cell
2. Cluster of cells > 95% mitochondria
3. High mitochondria %, low genes/cell

**Supplementary Table S6. The parameters set in Seurat analysis for single-cell RNA-seq data**

| | Min celle/gene | Min genes/cell | Max genes/cell | Max mito% | SCT features | Num PCs | Dims reduce | Cluster resulation | VST features |
|---|---|---|---|---|---|---|---|---|---|
| **Standard** | 3 | 200 | 5000 | 30 | 3000 | 50 | 20 | 0.8 | 2000 |
| **Adjusted-P6461_19617_met** | 3 | 200 | 5000 | 90 | 1000 | 100 | 100 | 0.8 | 2000 |
| **Adjusted-others** | 3 | 20 | 5000 | 90 | 2000 | 25 | 20 | 1 | 2000 |

Note: the adjusted others including P5847_18349_normal, P6342_19538B_normal and P6593_21332B_tumor

**Supplementary Table S7. The epithelial and non-epithelial markers for calculating module scores**

| Epithelial genes | | | | | | | |
|---|---|---|---|---|---|---|---|
| Gastric | Keratin | Tumor | Antrum | Intestine | Stem cells | Parietal | Neuroendocrine |
| MUC5AC | KRT7 | EPCAM | GAST | MUC2 | LGR5 | GIF | CHGA |
| PGC | KRT17 | TFF3 | PDX1 | | TROY | | GAST |
| MUC6 | KRT18 | CLDN4 | | | | | SST |
| TFF1 | KRT19 | | | | | | |
| TFF2 | | | | | | | |

| Non-epithelial genes | | | | | | |
|---|---|---|---|---|---|---|
| Fibrolast | Endothelium | Dendritic | Macrophage | Immune | B plasma | T cells |
| ACTA2 | VWF | CD83 | CD14 | PTRPC | CD19 | CD3D |
| DCN | PECAM1 | | FCGR3A | | CD79 | TRAC |
| SPARC | | | | | MS4A1 | TRAB |
| THY1 | | | | | IGHA | CD8A |
| | | | | | IGHC | NKG7 |
| | | | | | IGLC | GNLY |
| | | | | | | IL2RA |

**Supplementary Table S8: The most 3 significant pathways of each subclones across all patients**

| Patient | Pathway | ref.cluster | alt.cluster | ref.score | alt.score | adj.p.value |
|---|---|---|---|---|---|---|
| P5846 | ALLOGRAFT_REJECTION | normal | C1 | -0.707106781 | 0.707106781 | 8.11018E-13 |
| P5846 | APICAL_SURFACE | normal | C1 | -0.707106781 | 0.707106781 | 2.59163E-04 |
| P5846 | ESTROGEN_RESPONSE_EARLY | normal | C1 | 0.707106781 | -0.707106781 | 7.27443E-05 |
| P5846 | G2M_CHECKPOINT | normal | C1 | 0.707106781 | -0.707106781 | 2.01396E-04 |
| P5846 | INTERFERON_GAMMA_RESPONSE | normal | C1 | -0.707106781 | 0.707106781 | 1.23285E-06 |
| P5846 | MITOTIC_SPINDLE | normal | C1 | 0.707106781 | -0.707106781 | 6.02542E-05 |
| P5847 | ALLOGRAFT_REJECTION | C1C | C1A | 1.741809806 | -0.508174333 | 4.66081E-03 |
| P5847 | ALLOGRAFT_REJECTION | C1C | C2A | 1.741809806 | -0.724951511 | 1.28873E-03 |
| P5847 | ALLOGRAFT_REJECTION | C1C | C2B | 1.741809806 | -0.417011997 | 7.18148E-03 |
| P5847 | APICAL_SURFACE | C2B | C2A | 0.458731008 | -1.701235478 | 1.18598E-02 |
| P5847 | BILE_ACID_METABOLISM | C2A | C1A | -1.486827248 | 0.946056772 | 8.85771E-03 |
| P5847 | COAGULATION | C2A | C1A | -1.010793426 | -0.123519722 | 3.63195E-07 |
| P5847 | COAGULATION | C2B | C1A | -0.801598139 | -0.123519722 | 1.26699E-04 |
| P5847 | COAGULATION | C1C | C1A | 1.440503894 | -0.123519722 | 1.02112E-02 |
| P5847 | COAGULATION | C1B | C2A | 0.495407391 | -1.010793426 | 5.07618E-10 |
| P5847 | COAGULATION | C1C | C2A | 1.440503894 | -1.010793426 | 3.56059E-06 |
| P5847 | COAGULATION | C2B | C1B | -0.801598139 | 0.495407391 | 7.91326E-08 |
| P5847 | COAGULATION | C1C | C2B | 1.440503894 | -0.801598139 | 2.89705E-05 |
| P5847 | HEME_METABOLISM | C1B | C2A | 0.826359034 | -1.167962207 | 9.90097E-03 |
| P5847 | INFLAMMATORY_RESPONSE | C1C | C2A | 1.641026062 | -0.600938196 | 4.66152E-02 |
| P5847 | INFLAMMATORY_RESPONSE | C2B | C1B | -0.780731376 | 0.259379832 | 4.84412E-02 |
| P5847 | INFLAMMATORY_RESPONSE | C1C | C2B | 1.641026062 | -0.780731376 | 2.37713E-02 |
| P5847 | KRAS_SIGNALING_DN | C2A | C1A | 1.222768278 | -0.535357593 | 4.63715E-02 |
| P5847 | KRAS_SIGNALING_DN | C1B | C2A | -1.299097337 | 1.222768278 | 4.24218E-02 |
| P5847 | KRAS_SIGNALING_UP | C1B | C2A | 0.848099504 | -0.943318067 | 3.78739E-02 |
| P5847 | KRAS_SIGNALING_UP | C2B | C1B | -0.985882489 | 0.848099504 | 2.70172E-02 |
| P5847 | NOTCH_SIGNALING | C1C | C1A | 1.727562172 | -0.484150889 | 4.41073E-02 |
| P5847 | NOTCH_SIGNALING | C1C | C2A | 1.727562172 | -0.802420489 | 1.29623E-02 |
| P5847 | SPERMATOGENESIS | C1C | C2A | -1.757183218 | 0.75887645 | 2.34642E-02 |
| P5847 | TNFA_SIGNALING_VIA_NFKB | C1C | C2A | 1.717598088 | -0.820189641 | 4.68027E-03 |
| P5847 | TNFA_SIGNALING_VIA_NFKB | C1C | C2B | 1.717598088 | -0.553125176 | 1.55543E-02 |
| P5931 | ALLOGRAFT_REJECTION | C3 | normal | 0.432299048 | -1.849732378 | 4.76256E-11 |
| P5931 | ALLOGRAFT_REJECTION | C4B | normal | 0.369195484 | -1.849732378 | 4.76256E-11 |

| | | | | | | |
|---|---|---|---|---|---|---|
| P5931 | ALLOGRAFT_REJECTION | C1 | normal | 0.265580753 | -1.849732378 | 4.76256E-11 |
| P5931 | ALLOGRAFT_REJECTION | C2 | normal | 1.05408535 | -1.849732378 | 4.76256E-11 |
| P5931 | ALLOGRAFT_REJECTION | C4A | normal | -0.271428256 | -1.849732378 | 2.95129E-08 |
| P5931 | ALLOGRAFT_REJECTION | C2 | C3 | 1.05408535 | 0.432299048 | 8.38319E-06 |
| P5931 | ALLOGRAFT_REJECTION | C2 | C4B | 1.05408535 | 0.369195484 | 1.39044E-05 |
| P5931 | ALLOGRAFT_REJECTION | C2 | C1 | 1.05408535 | 0.265580753 | 4.84706E-11 |
| P5931 | ALLOGRAFT_REJECTION | C4A | C2 | -0.271428256 | 1.05408535 | 1.60392E-06 |
| P5931 | BILE_ACID_METABOLISM | C3 | normal | -0.413063377 | -1.593770016 | 7.78133E-03 |
| P5931 | BILE_ACID_METABOLISM | C4B | normal | 1.10723638 | -1.593770016 | 5.69744E-11 |
| P5931 | BILE_ACID_METABOLISM | C1 | normal | 0.741702139 | -1.593770016 | 4.81946E-11 |
| P5931 | BILE_ACID_METABOLISM | C2 | normal | 0.579305271 | -1.593770016 | 1.55517E-09 |
| P5931 | BILE_ACID_METABOLISM | C4B | C3 | 1.10723638 | -0.413063377 | 6.41858E-05 |
| P5931 | BILE_ACID_METABOLISM | C1 | C3 | 0.741702139 | -0.413063377 | 7.19582E-05 |
| P5931 | BILE_ACID_METABOLISM | C2 | C3 | 0.579305271 | -0.413063377 | 6.95638E-03 |
| P5931 | EPITHELIAL_MESENCHYMAL_TRANSITION | C2 | normal | 1.723798161 | -0.864282585 | 7.88063E-04 |
| P5931 | EPITHELIAL_MESENCHYMAL_TRANSITION | C2 | C3 | 1.723798161 | -0.883237645 | 3.01689E-05 |
| P5931 | EPITHELIAL_MESENCHYMAL_TRANSITION | C2 | C1 | 1.723798161 | -0.347236118 | 1.03985E-04 |
| P5931 | G2M_CHECKPOINT | C3 | normal | 0.229972499 | -1.627614218 | 4.76804E-11 |
| P5931 | G2M_CHECKPOINT | C4B | normal | 0.192968246 | -1.627614218 | 4.76792E-11 |
| P5931 | G2M_CHECKPOINT | C1 | normal | 0.040192575 | -1.627614218 | 4.76812E-11 |
| P5931 | G2M_CHECKPOINT | C2 | normal | 1.471376136 | -1.627614218 | 4.76256E-11 |
| P5931 | G2M_CHECKPOINT | C4A | normal | -0.306895239 | -1.627614218 | 1.65454E-03 |
| P5931 | G2M_CHECKPOINT | C2 | C3 | 1.471376136 | 0.229972499 | 4.80788E-11 |
| P5931 | G2M_CHECKPOINT | C2 | C4B | 1.471376136 | 0.192968246 | 9.17855E-11 |
| P5931 | G2M_CHECKPOINT | C2 | C1 | 1.471376136 | 0.040192575 | 4.76816E-11 |
| P5931 | G2M_CHECKPOINT | C4A | C2 | -0.306895239 | 1.471376136 | 7.51592E-07 |
| P5931 | HEDGEHOG_SIGNALING | C3 | normal | -0.286130367 | 1.895186955 | 5.33847E-06 |
| P5931 | HEDGEHOG_SIGNALING | C4B | normal | -0.309199074 | 1.895186955 | 3.27831E-05 |
| P5931 | HEDGEHOG_SIGNALING | C1 | normal | 0.203185621 | 1.895186955 | 1.39065E-04 |
| P5931 | HEDGEHOG_SIGNALING | C2 | normal | -0.578405965 | 1.895186955 | 4.96214E-08 |
| P5931 | HEDGEHOG_SIGNALING | C4A | normal | -0.92463717 | 1.895186955 | 2.50814E-03 |
| P5931 | INFLAMMATORY_RESPONSE | C3 | normal | 0.050114209 | -1.544281421 | 1.11923E-03 |
| P5931 | INFLAMMATORY_RESPONSE | C4B | normal | 1.04174865 | -1.544281421 | 6.40027E-08 |
| P5931 | INFLAMMATORY_RESPONSE | C1 | normal | 0.833954241 | -1.544281421 | 7.75671E-10 |
| P5931 | INFLAMMATORY_RESPONSE | C2 | normal | 0.425340428 | -1.544281421 | 8.68101E-06 |
| P5931 | KRAS_SIGNALING_DN | C3 | normal | -0.221142041 | 1.800725075 | 4.76256E-11 |
| P5931 | KRAS_SIGNALING_DN | C4B | normal | -0.361955058 | 1.800725075 | 4.76256E-11 |

| | | | | | | |
|---|---|---|---|---|---|---|
| P5931 | KRAS_SIGNALING_DN | C1 | normal | -0.137334166 | 1.800725075 | 4.76256E-11 |
| P5931 | KRAS_SIGNALING_DN | C2 | normal | -1.238393247 | 1.800725075 | 4.76256E-11 |
| P5931 | KRAS_SIGNALING_DN | C4A | normal | 0.158099437 | 1.800725075 | 9.18596E-08 |
| P5931 | KRAS_SIGNALING_DN | C2 | C3 | -1.238393247 | -0.221142041 | 4.81472E-11 |
| P5931 | KRAS_SIGNALING_DN | C2 | C4B | -1.238393247 | -0.361955058 | 8.13345E-08 |
| P5931 | KRAS_SIGNALING_DN | C2 | C1 | -1.238393247 | -0.137334166 | 4.76851E-11 |
| P5931 | KRAS_SIGNALING_DN | C4A | C2 | 0.158099437 | -1.238393247 | 2.81028E-06 |
| P5931 | KRAS_SIGNALING_UP | C4B | C3 | 1.625456689 | -1.012088212 | 5.56375E-03 |
| P5931 | KRAS_SIGNALING_UP | C1 | C3 | 0.795589013 | -1.012088212 | 1.90832E-02 |
| P5931 | KRAS_SIGNALING_UP | C2 | C4B | -0.527870715 | 1.625456689 | 3.58925E-02 |
| P5931 | MYC_TARGETS_V2 | C3 | normal | 0.447520638 | -1.833057466 | 4.76256E-11 |
| P5931 | MYC_TARGETS_V2 | C4B | normal | 0.282194433 | -1.833057466 | 4.76256E-11 |
| P5931 | MYC_TARGETS_V2 | C1 | normal | -0.082833802 | -1.833057466 | 4.76256E-11 |
| P5931 | MYC_TARGETS_V2 | C2 | normal | 1.163010673 | -1.833057466 | 4.76256E-11 |
| P5931 | MYC_TARGETS_V2 | C4A | normal | 0.023165524 | -1.833057466 | 2.19935E-09 |
| P5931 | MYC_TARGETS_V2 | C1 | C3 | -0.082833802 | 0.447520638 | 1.57663E-04 |
| P5931 | MYC_TARGETS_V2 | C2 | C3 | 1.163010673 | 0.447520638 | 2.49517E-06 |
| P5931 | MYC_TARGETS_V2 | C2 | C4B | 1.163010673 | 0.282194433 | 1.52055E-07 |
| P5931 | MYC_TARGETS_V2 | C2 | C1 | 1.163010673 | -0.082833802 | 4.76752E-11 |
| P5931 | MYC_TARGETS_V2 | C4A | C2 | 0.023165524 | 1.163010673 | 4.57858E-04 |
| P5931 | PANCREAS_BETA_CELLS | C3 | normal | 0.746381194 | -1.958317602 | 4.90220E-08 |
| P5931 | PANCREAS_BETA_CELLS | C4B | normal | 0.131530746 | -1.958317602 | 3.62328E-04 |
| P5931 | PANCREAS_BETA_CELLS | C1 | normal | 0.023730376 | -1.958317602 | 1.68738E-05 |
| P5931 | PANCREAS_BETA_CELLS | C2 | normal | 0.402316846 | -1.958317602 | 1.69727E-06 |
| P5931 | PANCREAS_BETA_CELLS | C4A | normal | 0.654358441 | -1.958317602 | 1.43593E-02 |
| P5931 | SPERMATOGENESIS | C4B | normal | -0.745919918 | 1.654467909 | 4.78521E-02 |
| P5931 | SPERMATOGENESIS | C1 | normal | -0.792483163 | 1.654467909 | 5.03859E-03 |
| P5931 | SPERMATOGENESIS | C2 | normal | -0.4887088 | 1.654467909 | 4.93851E-02 |
| P5931 | WNT_BETA_CATENIN_SIGNALING | C2 | normal | 1.791979542 | -0.866315557 | 8.69143E-05 |
| P5931 | WNT_BETA_CATENIN_SIGNALING | C2 | C3 | 1.791979542 | -0.109010374 | 2.06543E-03 |
| P5931 | WNT_BETA_CATENIN_SIGNALING | C2 | C4B | 1.791979542 | -0.972579788 | 1.40067E-05 |
| P5931 | WNT_BETA_CATENIN_SIGNALING | C2 | C1 | 1.791979542 | -0.106909145 | 9.70580E-05 |
| P6342 | ANGIOGENESIS | C1B | normal | -0.775783044 | -0.130550234 | 3.13917E-02 |
| P6342 | ANGIOGENESIS | C1A | normal | -1.048962933 | -0.130550234 | 6.83156E-04 |
| P6342 | ANGIOGENESIS | C3B | normal | 1.407300544 | -0.130550234 | 1.91166E-02 |
| P6342 | ANGIOGENESIS | C3A | C1B | 0.547995667 | -0.775783044 | 2.59522E-05 |
| P6342 | ANGIOGENESIS | C3B | C1B | 1.407300544 | -0.775783044 | 2.00088E-04 |

| | | | | | | |
|---|---|---|---|---|---|---|
| P6342 | ANGIOGENESIS | C3A | C1A | 0.547995667 | -1.048962933 | 3.11042E-07 |
| P6342 | ANGIOGENESIS | C3B | C1A | 1.407300544 | -1.048962933 | 1.96460E-05 |
| P6342 | BILE_ACID_METABOLISM | C1A | normal | 1.637954726 | -0.438214459 | 3.32961E-04 |
| P6342 | BILE_ACID_METABOLISM | C1A | C1B | 1.637954726 | 0.056978045 | 2.73205E-02 |
| P6342 | BILE_ACID_METABOLISM | C3A | C1A | -1.035885503 | 1.637954726 | 1.75086E-04 |
| P6342 | E2F_TARGETS | C1B | normal | 0.053720856 | -1.501913223 | 5.98453E-07 |
| P6342 | E2F_TARGETS | C1A | normal | 0.857439408 | -1.501913223 | 3.82472E-13 |
| P6342 | E2F_TARGETS | C3A | normal | 0.940230677 | -1.501913223 | 1.17034E-11 |
| P6342 | EPITHELIAL_MESENCHYMAL_TRANSITION | C3A | normal | 1.037396396 | -0.808077978 | 4.33986E-13 |
| P6342 | EPITHELIAL_MESENCHYMAL_TRANSITION | C3B | normal | 1.141498186 | -0.808077978 | 1.26317E-04 |
| P6342 | EPITHELIAL_MESENCHYMAL_TRANSITION | C3A | C1B | 1.037396396 | -0.804218046 | 2.72748E-12 |
| P6342 | EPITHELIAL_MESENCHYMAL_TRANSITION | C3B | C1B | 1.141498186 | -0.804218046 | 1.79267E-04 |
| P6342 | EPITHELIAL_MESENCHYMAL_TRANSITION | C3A | C1A | 1.037396396 | -0.566598558 | 3.80174E-09 |
| P6342 | EPITHELIAL_MESENCHYMAL_TRANSITION | C3B | C1A | 1.141498186 | -0.566598558 | 1.73308E-03 |
| P6342 | ESTROGEN_RESPONSE_LATE | C1B | normal | -0.422679117 | -1.459318533 | 1.64524E-12 |
| P6342 | ESTROGEN_RESPONSE_LATE | C1A | normal | 0.14520665 | -1.459318533 | 3.09530E-13 |
| P6342 | ESTROGEN_RESPONSE_LATE | C3A | normal | 0.583417146 | -1.459318533 | 2.92766E-13 |
| P6342 | ESTROGEN_RESPONSE_LATE | C3B | normal | 1.153373854 | -1.459318533 | 3.46168E-13 |
| P6342 | ESTROGEN_RESPONSE_LATE | C1A | C1B | 0.14520665 | -0.422679117 | 2.18169E-03 |
| P6342 | ESTROGEN_RESPONSE_LATE | C3A | C1B | 0.583417146 | -0.422679117 | 9.67273E-08 |
| P6342 | ESTROGEN_RESPONSE_LATE | C3B | C1B | 1.153373854 | -0.422679117 | 7.59017E-06 |
| P6342 | ESTROGEN_RESPONSE_LATE | C3B | C1A | 1.153373854 | 0.14520665 | 1.40345E-02 |
| P6342 | G2M_CHECKPOINT | C1B | normal | -0.34539089 | -1.402564855 | 4.04762E-03 |
| P6342 | G2M_CHECKPOINT | C1A | normal | 0.882711287 | -1.402564855 | 3.80418E-12 |
| P6342 | G2M_CHECKPOINT | C3A | normal | 1.049069208 | -1.402564855 | 1.10214E-10 |
| P6342 | G2M_CHECKPOINT | C1A | C1B | 0.882711287 | -0.34539089 | 2.28288E-03 |
| P6342 | G2M_CHECKPOINT | C3A | C1B | 1.049069208 | -0.34539089 | 2.17044E-03 |
| P6342 | HEDGEHOG_SIGNALING | C1B | normal | -0.087556436 | 1.738248287 | 3.40283E-13 |
| P6342 | HEDGEHOG_SIGNALING | C1A | normal | -0.733550256 | 1.738248287 | 2.92766E-13 |
| P6342 | HEDGEHOG_SIGNALING | C3A | normal | -0.380607881 | 1.738248287 | 3.44946E-13 |
| P6342 | HEDGEHOG_SIGNALING | C3B | normal | -0.536533713 | 1.738248287 | 2.30513E-06 |
| P6342 | HEDGEHOG_SIGNALING | C1A | C1B | -0.733550256 | -0.087556436 | 2.34262E-02 |
| P6342 | HEME_METABOLISM | C1B | normal | 0.087447621 | -1.289030563 | 3.39839E-13 |
| P6342 | HEME_METABOLISM | C1A | normal | 1.502017616 | -1.289030563 | 2.92766E-13 |
| P6342 | HEME_METABOLISM | C3A | normal | -0.02863258 | -1.289030563 | 4.56080E-13 |
| P6342 | HEME_METABOLISM | C3B | normal | -0.271802095 | -1.289030563 | 7.89029E-03 |
| P6342 | HEME_METABOLISM | C1A | C1B | 1.502017616 | 0.087447621 | 3.41505E-13 |

| | | | | | | |
|---|---|---|---|---|---|---|
| P6342 | HEME_METABOLISM | C3A | C1A | -0.02863258 | 1.502017616 | 3.34399E-13 |
| P6342 | HEME_METABOLISM | C3B | C1A | -0.271802095 | 1.502017616 | 1.69996E-07 |
| P6342 | INTERFERON_ALPHA_RESPONSE | C1B | normal | 0.609832014 | -1.038840905 | 2.92766E-13 |
| P6342 | INTERFERON_ALPHA_RESPONSE | C1A | normal | 1.43072978 | -1.038840905 | 2.92766E-13 |
| P6342 | INTERFERON_ALPHA_RESPONSE | C3A | normal | -0.490750657 | -1.038840905 | 5.48817E-03 |
| P6342 | INTERFERON_ALPHA_RESPONSE | C1A | C1B | 1.43072978 | 0.609832014 | 4.45722E-07 |
| P6342 | INTERFERON_ALPHA_RESPONSE | C3A | C1B | -0.490750657 | 0.609832014 | 1.00212E-09 |
| P6342 | INTERFERON_ALPHA_RESPONSE | C3B | C1B | -0.510970232 | 0.609832014 | 2.53251E-03 |
| P6342 | INTERFERON_ALPHA_RESPONSE | C3A | C1A | -0.490750657 | 1.43072978 | 3.12750E-13 |
| P6342 | INTERFERON_ALPHA_RESPONSE | C3B | C1A | -0.510970232 | 1.43072978 | 4.72451E-09 |
| P6342 | INTERFERON_GAMMA_RESPONSE | C1B | normal | 0.500796014 | -1.043924466 | 3.09974E-13 |
| P6342 | INTERFERON_GAMMA_RESPONSE | C1A | normal | 1.483649338 | -1.043924466 | 2.92766E-13 |
| P6342 | INTERFERON_GAMMA_RESPONSE | C3A | normal | -0.381041798 | -1.043924466 | 5.48721E-04 |
| P6342 | INTERFERON_GAMMA_RESPONSE | C1A | C1B | 1.483649338 | 0.500796014 | 2.17263E-09 |
| P6342 | INTERFERON_GAMMA_RESPONSE | C3A | C1B | -0.381041798 | 0.500796014 | 4.35711E-06 |
| P6342 | INTERFERON_GAMMA_RESPONSE | C3B | C1B | -0.559479089 | 0.500796014 | 7.32294E-03 |
| P6342 | INTERFERON_GAMMA_RESPONSE | C3A | C1A | -0.381041798 | 1.483649338 | 3.39395E-13 |
| P6342 | INTERFERON_GAMMA_RESPONSE | C3B | C1A | -0.559479089 | 1.483649338 | 1.97376E-09 |
| P6342 | KRAS_SIGNALING_DN | C1B | normal | -0.341222058 | 1.611386338 | 2.92766E-13 |
| P6342 | KRAS_SIGNALING_DN | C1A | normal | -1.115349293 | 1.611386338 | 2.92766E-13 |
| P6342 | KRAS_SIGNALING_DN | C3A | normal | -0.201933979 | 1.611386338 | 2.92766E-13 |
| P6342 | KRAS_SIGNALING_DN | C3B | normal | 0.047118991 | 1.611386338 | 4.17088E-09 |
| P6342 | KRAS_SIGNALING_DN | C1A | C1B | -1.115349293 | -0.341222058 | 3.68983E-09 |
| P6342 | KRAS_SIGNALING_DN | C3A | C1A | -0.201933979 | -1.115349293 | 1.62805E-09 |
| P6342 | KRAS_SIGNALING_DN | C3B | C1A | 0.047118991 | -1.115349293 | 5.11681E-05 |
| P6342 | REACTIVE_OXIGEN_SPECIES_PATHWAY | C1B | normal | 0.527554738 | -1.484415713 | 2.92766E-13 |
| P6342 | REACTIVE_OXIGEN_SPECIES_PATHWAY | C1A | normal | 1.217936762 | -1.484415713 | 2.92766E-13 |
| P6342 | REACTIVE_OXIGEN_SPECIES_PATHWAY | C3A | normal | -0.111227932 | -1.484415713 | 3.40061E-13 |
| P6342 | REACTIVE_OXIGEN_SPECIES_PATHWAY | C3B | normal | -0.149847857 | -1.484415713 | 1.77590E-05 |
| P6342 | REACTIVE_OXIGEN_SPECIES_PATHWAY | C1A | C1B | 1.217936762 | 0.527554738 | 5.56079E-06 |
| P6342 | REACTIVE_OXIGEN_SPECIES_PATHWAY | C3A | C1B | -0.111227932 | 0.527554738 | 3.97225E-04 |
| P6342 | REACTIVE_OXIGEN_SPECIES_PATHWAY | C3A | C1A | -0.111227932 | 1.217936762 | 3.46057E-13 |
| P6342 | REACTIVE_OXIGEN_SPECIES_PATHWAY | C3B | C1A | -0.149847857 | 1.217936762 | 1.70592E-05 |
| P6342 | SPERMATOGENESIS | C1B | normal | -0.357330678 | 1.314270271 | 3.09475E-12 |
| P6342 | SPERMATOGENESIS | C1A | normal | -1.376552568 | 1.314270271 | 2.95874E-13 |
| P6342 | SPERMATOGENESIS | C3A | normal | 0.494476421 | 1.314270271 | 2.00589E-02 |
| P6342 | SPERMATOGENESIS | C1A | C1B | -1.376552568 | -0.357330678 | 5.21107E-04 |

| | | | | | | |
|---|---|---|---|---|---|---|
| P6342 | SPERMATOGENESIS | C3A | C1B | 0.494476421 | -0.357330678 | 2.36019E-02 |
| P6342 | SPERMATOGENESIS | C3A | C1A | 0.494476421 | -1.376552568 | 1.84538E-09 |
| P6342 | TNFA_SIGNALING_VIA_NFKB | C1A | normal | 1.651114339 | -0.776110284 | 3.41061E-13 |
| P6342 | TNFA_SIGNALING_VIA_NFKB | C3A | normal | 0.221602291 | -0.776110284 | 6.75908E-03 |
| P6342 | TNFA_SIGNALING_VIA_NFKB | C1A | C1B | 1.651114339 | -0.4456047 | 9.66671E-13 |
| P6342 | TNFA_SIGNALING_VIA_NFKB | C3A | C1A | 0.221602291 | 1.651114339 | 8.09557E-05 |
| P6342 | TNFA_SIGNALING_VIA_NFKB | C3B | C1A | -0.651001646 | 1.651114339 | 5.90811E-04 |
| P5915 | ADIPOGENESIS | C1 | normal | -0.707106781 | 0.707106781 | 1.02699E-03 |
| P5915 | ALLOGRAFT_REJECTION | C1 | normal | -0.707106781 | 0.707106781 | 2.17077E-02 |
| P5915 | E2F_TARGETS | C1 | normal | -0.707106781 | 0.707106781 | 1.79601E-02 |
| P5915 | KRAS_SIGNALING_DN | C1 | normal | 0.707106781 | -0.707106781 | 2.54881E-03 |
| P5915 | P53_PATHWAY | C1 | normal | -0.707106781 | 0.707106781 | 5.97078E-13 |
| P5915 | SPERMATOGENESIS | C1 | normal | -0.707106781 | 0.707106781 | 6.42501E-04 |
| P5915 | XENOBIOTIC_METABOLISM | C1 | normal | -0.707106781 | 0.707106781 | 5.23109E-09 |
| P6198 | ADIPOGENESIS | C1 | normal | 0.761922464 | -1.4386513 | 3.30360E-02 |
| P6198 | ADIPOGENESIS | C3 | C1 | 0.092625089 | 0.761922464 | 7.66188E-03 |
| P6198 | APICAL_SURFACE | C2 | C1 | 0.144206941 | -0.765879814 | 1.59343E-02 |
| P6198 | ESTROGEN_RESPONSE_EARLY | C2 | C1 | 0.1446752 | 0.948413077 | 6.28682E-06 |
| P6198 | FATTY_ACID_METABOLISM | C3 | C1 | 0.111770859 | 0.741770088 | 3.50348E-02 |
| P6198 | HEME_METABOLISM | C2 | C1 | 0.435625139 | 0.997165997 | 2.26550E-03 |
| P6198 | HEME_METABOLISM | C3 | C1 | -0.088125924 | 0.997165997 | 2.29937E-04 |
| P6198 | HYPOXIA | C2 | C1 | 0.224905486 | 1.115407377 | 4.78468E-03 |
| P6198 | HYPOXIA | C3 | C1 | -0.034914501 | 1.115407377 | 4.75355E-02 |
| P6198 | PI3K_AKT_MTOR_SIGNALING | C1 | normal | 0.830586482 | -1.454284131 | 4.35155E-02 |
| P6198 | PI3K_AKT_MTOR_SIGNALING | C2 | C1 | 0.293350417 | 0.830586482 | 4.42962E-04 |
| P6198 | PROTEIN_SECRETION | C1 | normal | 0.730999683 | -1.474452463 | 3.61352E-02 |
| P6198 | TNFA_SIGNALING_VIA_NFKB | C2 | C1 | 0.28021661 | 1.217903836 | 2.17916E-02 |
| P6198 | TNFA_SIGNALING_VIA_NFKB | C3 | C1 | -0.35156402 | 1.217903836 | 2.06691E-02 |
| P6198 | UV_RESPONSE_DN | C3 | C1 | -0.108249539 | 0.847893041 | 3.97838E-02 |
| P6335 | ALLOGRAFT_REJECTION | C2 | normal | -1.367956429 | 1.000913472 | 4.86370E-04 |
| P6335 | ALLOGRAFT_REJECTION | C2 | C1 | -1.367956429 | 0.011029212 | 5.71361E-03 |
| P6335 | ALLOGRAFT_REJECTION | C2 | C3 | -1.367956429 | 0.356013745 | 5.11507E-03 |
| P6335 | APICAL_SURFACE | C1 | normal | -0.665337182 | 1.46936041 | 1.16402E-08 |
| P6335 | APICAL_SURFACE | C3 | normal | -0.208259525 | 1.46936041 | 1.91218E-04 |
| P6335 | APICAL_SURFACE | C2 | normal | -0.595763703 | 1.46936041 | 6.81475E-07 |
| P6335 | IL6_JAK_STAT3_SIGNALING | C2 | normal | -1.308366042 | 1.103892219 | 2.04500E-03 |
| P6335 | IL6_JAK_STAT3_SIGNALING | C2 | C1 | -1.308366042 | -0.053800739 | 3.91230E-02 |

| | | | | | | |
|---|---|---|---|---|---|---|
| P6335 | IL6_JAK_STAT3_SIGNALING | C2 | C3 | -1.308366042 | 0.258274561 | 3.67018E-02 |
| P6335 | INFLAMMATORY_RESPONSE | C1 | normal | -0.462820059 | 1.435419869 | 3.11125E-06 |
| P6335 | INFLAMMATORY_RESPONSE | C3 | normal | -0.131059004 | 1.435419869 | 1.59381E-03 |
| P6335 | INFLAMMATORY_RESPONSE | C2 | normal | -0.841540806 | 1.435419869 | 2.84601E-07 |
| P6335 | KRAS_SIGNALING_DN | C1 | normal | -1.455171244 | 0.75788667 | 1.09141E-03 |
| P6335 | KRAS_SIGNALING_DN | C3 | C1 | 0.168390303 | -1.455171244 | 5.67510E-03 |
| P6335 | KRAS_SIGNALING_DN | C2 | C1 | 0.528894272 | -1.455171244 | 8.52992E-05 |
| P6335 | MYC_TARGETS_V1 | C3 | C1 | -0.721951312 | 1.472462803 | 4.35128E-03 |
| P6335 | MYC_TARGETS_V1 | C2 | C1 | -0.495683832 | 1.472462803 | 5.89392E-03 |
| P6335 | OXIDATIVE_PHOSPHORYLATION | C3 | C1 | -0.468347912 | 1.462834425 | 6.45741E-03 |
| P6335 | OXIDATIVE_PHOSPHORYLATION | C2 | C1 | -0.767672515 | 1.462834425 | 2.91508E-04 |
| P6335 | PI3K_AKT_MTOR_SIGNALING | C3 | C1 | -0.779009296 | 1.455513341 | 4.68693E-04 |
| P6335 | PI3K_AKT_MTOR_SIGNALING | C2 | C1 | -0.489573816 | 1.455513341 | 1.10445E-03 |
| P6335 | SPERMATOGENESIS | C1 | normal | -1.215007784 | 0.524513553 | 1.14217E-02 |
| P6335 | SPERMATOGENESIS | C2 | C1 | 1.055968747 | -1.215007784 | 1.43165E-06 |
| P6335 | SPERMATOGENESIS | C2 | C3 | 1.055968747 | -0.365474516 | 4.14189E-02 |
| P6335 | UV_RESPONSE_DN | C1 | normal | -0.205216452 | 1.461488697 | 2.18789E-03 |
| P6335 | UV_RESPONSE_DN | C3 | normal | -0.500020906 | 1.461488697 | 1.54971E-03 |
| P6335 | UV_RESPONSE_DN | C2 | normal | -0.75625134 | 1.461488697 | 1.09447E-04 |
| P6335 | WNT_BETA_CATENIN_SIGNALING | C1 | normal | 1.338296384 | -0.932308815 | 7.93882E-04 |
| P6335 | WNT_BETA_CATENIN_SIGNALING | C3 | C1 | -0.561706838 | 1.338296384 | 7.73981E-04 |
| P6335 | WNT_BETA_CATENIN_SIGNALING | C2 | C1 | 0.155719269 | 1.338296384 | 4.71421E-02 |
| P6461 | ALLOGRAFT_REJECTION | C6 | C2 | -0.631903704 | 0.770870786 | 1.75482E-08 |
| P6461 | ALLOGRAFT_REJECTION | C3B | C2 | -1.189728332 | 0.770870786 | 2.87738E-05 |
| P6461 | ALLOGRAFT_REJECTION | C4 | C2 | -0.621690441 | 0.770870786 | 7.86527E-03 |
| P6461 | ALLOGRAFT_REJECTION | C1A | C2 | -1.288716792 | 0.770870786 | 6.17153E-03 |
| P6461 | ALLOGRAFT_REJECTION | C5 | C6 | 0.46690302 | -0.631903704 | 3.03224E-04 |
| P6461 | ALLOGRAFT_REJECTION | met | C6 | 1.726227977 | -0.631903704 | 2.38666E-03 |
| P6461 | ALLOGRAFT_REJECTION | C3B | C1B | -1.189728332 | 0.224472174 | 3.50176E-02 |
| P6461 | ALLOGRAFT_REJECTION | C3B | C5 | -1.189728332 | 0.46690302 | 1.76687E-03 |
| P6461 | ALLOGRAFT_REJECTION | C1A | C5 | -1.288716792 | 0.46690302 | 4.71965E-02 |
| P6461 | ALLOGRAFT_REJECTION | met | C3B | 1.726227977 | -1.189728332 | 5.96169E-04 |
| P6461 | ALLOGRAFT_REJECTION | met | C4 | 1.726227977 | -0.621690441 | 1.33881E-02 |
| P6461 | ALLOGRAFT_REJECTION | met | C1A | 1.726227977 | -1.288716792 | 3.60453E-03 |
| P6461 | APICAL_JUNCTION | C1B | C2 | 0.267437402 | -0.649029028 | 2.18239E-02 |
| P6461 | APICAL_SURFACE | C1B | C2 | 0.891169576 | -0.036195457 | 1.88989E-03 |
| P6461 | APICAL_SURFACE | C3A | C1B | -2.543910666 | 0.891169576 | 3.39033E-02 |

| | | | | | | |
|---|---|---|---|---|---|---|
| P6461 | DNA_REPAIR | met | C2 | 2.602379981 | -0.309210908 | 0.00000E+00 |
| P6461 | DNA_REPAIR | met | C6 | 2.602379981 | -0.4840864 | 0.00000E+00 |
| P6461 | DNA_REPAIR | met | C1B | 2.602379981 | -0.316843388 | 0.00000E+00 |
| P6461 | DNA_REPAIR | met | C3A | 2.602379981 | 0.162076589 | 1.09663E-06 |
| P6461 | DNA_REPAIR | met | C5 | 2.602379981 | -0.25777176 | 0.00000E+00 |
| P6461 | DNA_REPAIR | met | C3B | 2.602379981 | -0.428865637 | 0.00000E+00 |
| P6461 | DNA_REPAIR | met | C4 | 2.602379981 | -0.321919469 | 0.00000E+00 |
| P6461 | DNA_REPAIR | met | C1A | 2.602379981 | -0.645759006 | 0.00000E+00 |
| P6461 | E2F_TARGETS | met | C2 | 2.597927227 | -0.455003204 | 0.00000E+00 |
| P6461 | E2F_TARGETS | met | C6 | 2.597927227 | -0.303330481 | 0.00000E+00 |
| P6461 | E2F_TARGETS | met | C1B | 2.597927227 | -0.316020453 | 0.00000E+00 |
| P6461 | E2F_TARGETS | met | C3A | 2.597927227 | 0.048111313 | 1.06442E-02 |
| P6461 | E2F_TARGETS | met | C5 | 2.597927227 | -0.33454081 | 0.00000E+00 |
| P6461 | E2F_TARGETS | met | C3B | 2.597927227 | -0.02300054 | 6.39366E-09 |
| P6461 | E2F_TARGETS | met | C4 | 2.597927227 | -0.611251515 | 0.00000E+00 |
| P6461 | E2F_TARGETS | met | C1A | 2.597927227 | -0.602891538 | 4.70553E-10 |
| P6461 | HEME_METABOLISM | met | C2 | 2.625787458 | -0.284670846 | 0.00000E+00 |
| P6461 | HEME_METABOLISM | met | C6 | 2.625787458 | -0.375083719 | 0.00000E+00 |
| P6461 | HEME_METABOLISM | met | C1B | 2.625787458 | -0.200502933 | 0.00000E+00 |
| P6461 | HEME_METABOLISM | met | C3A | 2.625787458 | -0.112192825 | 4.21251E-03 |
| P6461 | HEME_METABOLISM | met | C5 | 2.625787458 | -0.246774713 | 0.00000E+00 |
| P6461 | HEME_METABOLISM | met | C3B | 2.625787458 | -0.237840579 | 1.43613E-10 |
| P6461 | HEME_METABOLISM | met | C4 | 2.625787458 | -0.461265415 | 2.66787E-13 |
| P6461 | HEME_METABOLISM | met | C1A | 2.625787458 | -0.707456429 | 7.99112E-11 |
| P6461 | IL2_STAT5_SIGNALING | C5 | C2 | 0.081920537 | -0.722651889 | 2.44070E-02 |
| P6461 | IL6_JAK_STAT3_SIGNALING | C6 | C2 | 1.31560099 | 0.292537221 | 4.99415E-02 |
| P6461 | IL6_JAK_STAT3_SIGNALING | met | C6 | -2.212505551 | 1.31560099 | 1.80107E-03 |
| P6461 | INFLAMMATORY_RESPONSE | met | C2 | -2.625477023 | 0.148171278 | 0.00000E+00 |
| P6461 | INFLAMMATORY_RESPONSE | met | C6 | -2.625477023 | 0.308480394 | 0.00000E+00 |
| P6461 | INFLAMMATORY_RESPONSE | met | C1B | -2.625477023 | 0.436813457 | 0.00000E+00 |
| P6461 | INFLAMMATORY_RESPONSE | met | C3A | -2.625477023 | 0.053746699 | 7.68887E-05 |
| P6461 | INFLAMMATORY_RESPONSE | met | C5 | -2.625477023 | 0.387420656 | 0.00000E+00 |
| P6461 | INFLAMMATORY_RESPONSE | met | C3B | -2.625477023 | 0.518293299 | 0.00000E+00 |
| P6461 | INFLAMMATORY_RESPONSE | met | C4 | -2.625477023 | 0.187629702 | 0.00000E+00 |
| P6461 | INFLAMMATORY_RESPONSE | met | C1A | -2.625477023 | 0.584921538 | 0.00000E+00 |
| P6461 | KRAS_SIGNALING_DN | met | C2 | -2.548487317 | 0.300896422 | 0.00000E+00 |
| P6461 | KRAS_SIGNALING_DN | met | C6 | -2.548487317 | 0.444089888 | 0.00000E+00 |

| | | | | | | |
|---|---|---|---|---|---|---|
| P6461 | KRAS_SIGNALING_DN | met | C1B | -2.548487317 | 0.280599842 | 0.00000E+00 |
| P6461 | KRAS_SIGNALING_DN | met | C3A | -2.548487317 | -0.369472292 | 5.68308E-07 |
| P6461 | KRAS_SIGNALING_DN | met | C5 | -2.548487317 | 0.245335975 | 0.00000E+00 |
| P6461 | KRAS_SIGNALING_DN | met | C3B | -2.548487317 | 0.582983386 | 0.00000E+00 |
| P6461 | KRAS_SIGNALING_DN | met | C4 | -2.548487317 | 0.397775539 | 0.00000E+00 |
| P6461 | KRAS_SIGNALING_DN | met | C1A | -2.548487317 | 0.666278558 | 0.00000E+00 |
| P6461 | KRAS_SIGNALING_UP | C6 | C2 | 0.686919292 | 0.12909844 | 1.80792E-02 |
| P6461 | KRAS_SIGNALING_UP | C1B | C2 | 0.729427747 | 0.12909844 | 1.03236E-02 |
| P6461 | KRAS_SIGNALING_UP | met | C2 | -2.39919475 | 0.12909844 | 3.33938E-08 |
| P6461 | KRAS_SIGNALING_UP | met | C6 | -2.39919475 | 0.686919292 | 4.41763E-11 |
| P6461 | KRAS_SIGNALING_UP | met | C1B | -2.39919475 | 0.729427747 | 2.65403E-11 |
| P6461 | KRAS_SIGNALING_UP | met | C5 | -2.39919475 | 0.471007224 | 3.12925E-10 |
| P6461 | KRAS_SIGNALING_UP | met | C3B | -2.39919475 | 0.546559734 | 9.57522E-08 |
| P6461 | KRAS_SIGNALING_UP | met | C4 | -2.39919475 | 0.376903569 | 4.60241E-07 |
| P6461 | KRAS_SIGNALING_UP | met | C1A | -2.39919475 | 0.18486119 | 1.85750E-04 |
| P6461 | MYC_TARGETS_V2 | C3A | C2 | 1.776317333 | -0.458583203 | 1.84873E-02 |
| P6461 | MYC_TARGETS_V2 | met | C2 | 1.683100163 | -0.458583203 | 5.67470E-08 |
| P6461 | MYC_TARGETS_V2 | C3A | C6 | 1.776317333 | -0.35794736 | 3.55391E-02 |
| P6461 | MYC_TARGETS_V2 | met | C6 | 1.683100163 | -0.35794736 | 1.49495E-06 |
| P6461 | MYC_TARGETS_V2 | C3A | C1B | 1.776317333 | -0.307927064 | 4.52914E-02 |
| P6461 | MYC_TARGETS_V2 | met | C1B | 1.683100163 | -0.307927064 | 3.52522E-06 |
| P6461 | MYC_TARGETS_V2 | C5 | C3A | -0.334488194 | 1.776317333 | 3.59218E-02 |
| P6461 | MYC_TARGETS_V2 | C3B | C3A | -0.73809391 | 1.776317333 | 9.08255E-03 |
| P6461 | MYC_TARGETS_V2 | C4 | C3A | -0.38102609 | 1.776317333 | 4.72074E-02 |
| P6461 | MYC_TARGETS_V2 | C1A | C3A | -0.881351675 | 1.776317333 | 9.23249E-03 |
| P6461 | MYC_TARGETS_V2 | met | C5 | 1.683100163 | -0.334488194 | 7.71800E-07 |
| P6461 | MYC_TARGETS_V2 | met | C3B | 1.683100163 | -0.73809391 | 4.40250E-07 |
| P6461 | MYC_TARGETS_V2 | met | C4 | 1.683100163 | -0.38102609 | 2.91028E-05 |
| P6461 | MYC_TARGETS_V2 | met | C1A | 1.683100163 | -0.881351675 | 5.44199E-06 |
| P6461 | SPERMATOGENESIS | met | C2 | -2.508768111 | 0.029658288 | 6.74518E-06 |
| P6461 | SPERMATOGENESIS | met | C6 | -2.508768111 | 0.587555307 | 5.51922E-08 |
| P6461 | SPERMATOGENESIS | met | C1B | -2.508768111 | 0.180420671 | 5.92427E-06 |
| P6461 | SPERMATOGENESIS | met | C5 | -2.508768111 | -0.034068567 | 2.04701E-05 |
| P6461 | SPERMATOGENESIS | met | C3B | -2.508768111 | 0.808935558 | 4.99205E-07 |
| P6461 | SPERMATOGENESIS | met | C4 | -2.508768111 | 0.620546248 | 2.03485E-06 |
| P6461 | SPERMATOGENESIS | met | C1A | -2.508768111 | 0.502080649 | 2.49770E-04 |
| P6461 | TNFA_SIGNALING_VIA_NFKB | met | C3B | -2.276988456 | 1.227988136 | 3.98544E-02 |

| | | | | | | |
|---|---|---|---|---|---|---|
| P6593 | APICAL_SURFACE | met | C7 | 2.181683704 | -0.118616506 | 2.39070E-11 |
| P6593 | APICAL_SURFACE | met | C1 | 2.181683704 | 0.098014444 | 6.13306E-07 |
| P6593 | APICAL_SURFACE | met | C5 | 2.181683704 | -0.860580843 | 2.16852E-06 |
| P6593 | APICAL_SURFACE | met | C3 | 2.181683704 | -0.448101593 | 9.66487E-05 |
| P6593 | APICAL_SURFACE | met | C6 | 2.181683704 | -0.290690933 | 2.74700E-02 |
| P6593 | APICAL_SURFACE | met | C2 | 2.181683704 | -1.161927755 | 1.60310E-02 |
| P6593 | BILE_ACID_METABOLISM | met | C7 | -2.306898334 | 0.087479877 | 2.39918E-11 |
| P6593 | BILE_ACID_METABOLISM | met | C1 | -2.306898334 | 0.206628737 | 1.03397E-05 |
| P6593 | BILE_ACID_METABOLISM | met | C3 | -2.306898334 | 0.467654166 | 5.97465E-03 |
| P6593 | BILE_ACID_METABOLISM | met | C4 | -2.306898334 | 0.451410086 | 2.57312E-03 |
| P6593 | BILE_ACID_METABOLISM | met | C6 | -2.306898334 | 1.285652253 | 9.45236E-03 |
| P6593 | COMPLEMENT | met | C7 | 2.185958158 | -0.237513192 | 2.40019E-11 |
| P6593 | COMPLEMENT | met | C1 | 2.185958158 | -0.133372502 | 2.39407E-04 |
| P6593 | COMPLEMENT | met | C5 | 2.185958158 | -1.136583171 | 7.63722E-04 |
| P6593 | COMPLEMENT | met | C3 | 2.185958158 | -0.389771676 | 2.80907E-02 |
| P6593 | COMPLEMENT | met | C4 | 2.185958158 | -0.261784228 | 2.48405E-02 |
| P6593 | DNA_REPAIR | met | C7 | 2.661319586 | -0.228197096 | 2.39070E-11 |
| P6593 | DNA_REPAIR | met | C1 | 2.661319586 | -0.289447491 | 2.39070E-11 |
| P6593 | DNA_REPAIR | met | C8 | 2.661319586 | -0.401708968 | 2.40094E-11 |
| P6593 | DNA_REPAIR | met | C5 | 2.661319586 | -0.373042512 | 2.39070E-11 |
| P6593 | DNA_REPAIR | met | C3 | 2.661319586 | -0.385571423 | 2.39070E-11 |
| P6593 | DNA_REPAIR | met | C4 | 2.661319586 | -0.253436319 | 2.39070E-11 |
| P6593 | DNA_REPAIR | met | C6 | 2.661319586 | -0.335720304 | 2.39162E-11 |
| P6593 | DNA_REPAIR | met | C2 | 2.661319586 | -0.394195473 | 2.39864E-11 |
| P6593 | EPITHELIAL_MESENCHYMAL_TRANSITION | met | C7 | 2.220404728 | -0.317949215 | 2.39070E-11 |
| P6593 | EPITHELIAL_MESENCHYMAL_TRANSITION | met | C1 | 2.220404728 | -0.061031144 | 3.67602E-08 |
| P6593 | EPITHELIAL_MESENCHYMAL_TRANSITION | met | C5 | 2.220404728 | -0.691874794 | 1.00712E-05 |
| P6593 | EPITHELIAL_MESENCHYMAL_TRANSITION | met | C3 | 2.220404728 | -0.422796171 | 1.08725E-04 |
| P6593 | EPITHELIAL_MESENCHYMAL_TRANSITION | met | C4 | 2.220404728 | -0.722888183 | 1.21306E-06 |
| P6593 | EPITHELIAL_MESENCHYMAL_TRANSITION | met | C6 | 2.220404728 | -0.836307831 | 1.84405E-03 |
| P6593 | ESTROGEN_RESPONSE_EARLY | met | C7 | 2.542505728 | 0.207516106 | 2.39070E-11 |
| P6593 | ESTROGEN_RESPONSE_EARLY | met | C1 | 2.542505728 | -0.271039552 | 2.39947E-11 |
| P6593 | ESTROGEN_RESPONSE_EARLY | met | C8 | 2.542505728 | -0.405846269 | 1.72641E-02 |
| P6593 | ESTROGEN_RESPONSE_EARLY | met | C5 | 2.542505728 | -0.327208344 | 1.59062E-08 |
| P6593 | ESTROGEN_RESPONSE_EARLY | met | C3 | 2.542505728 | -0.003462127 | 1.07666E-06 |
| P6593 | ESTROGEN_RESPONSE_EARLY | met | C4 | 2.542505728 | -0.765153373 | 2.47873E-11 |
| P6593 | ESTROGEN_RESPONSE_EARLY | met | C6 | 2.542505728 | -0.693986973 | 5.11984E-06 |

| | | | | | | |
|---|---|---|---|---|---|---|
| P6593 | ESTROGEN_RESPONSE_EARLY | met | C2 | 2.542505728 | -0.283325198 | 1.07304E-02 |
| P6593 | G2M_CHECKPOINT | met | C7 | 2.655096468 | -0.314811571 | 2.39070E-11 |
| P6593 | G2M_CHECKPOINT | met | C1 | 2.655096468 | -0.389309426 | 2.39070E-11 |
| P6593 | G2M_CHECKPOINT | met | C8 | 2.655096468 | -0.20991301 | 5.85739E-04 |
| P6593 | G2M_CHECKPOINT | met | C5 | 2.655096468 | -0.327505073 | 2.40012E-11 |
| P6593 | G2M_CHECKPOINT | met | C3 | 2.655096468 | -0.336664377 | 2.39994E-11 |
| P6593 | G2M_CHECKPOINT | met | C4 | 2.655096468 | -0.516747701 | 2.39878E-11 |
| P6593 | G2M_CHECKPOINT | met | C6 | 2.655096468 | -0.205175814 | 7.78772E-08 |
| P6593 | G2M_CHECKPOINT | met | C2 | 2.655096468 | -0.354969496 | 3.47600E-05 |
| P6593 | HEDGEHOG_SIGNALING | met | C7 | -2.543357964 | 0.016041157 | 2.39070E-11 |
| P6593 | HEDGEHOG_SIGNALING | met | C1 | -2.543357964 | 0.054119384 | 2.40061E-11 |
| P6593 | HEDGEHOG_SIGNALING | met | C8 | -2.543357964 | 0.984356125 | 2.27839E-03 |
| P6593 | HEDGEHOG_SIGNALING | met | C5 | -2.543357964 | 0.021501198 | 2.30648E-06 |
| P6593 | HEDGEHOG_SIGNALING | met | C3 | -2.543357964 | 0.370609907 | 3.07610E-08 |
| P6593 | HEDGEHOG_SIGNALING | met | C4 | -2.543357964 | 0.445571355 | 8.20131E-10 |
| P6593 | HEDGEHOG_SIGNALING | met | C6 | -2.543357964 | 0.399184134 | 1.24405E-04 |
| P6593 | HEDGEHOG_SIGNALING | met | C2 | -2.543357964 | 0.251974703 | 1.81180E-02 |
| P6593 | IL2_STAT5_SIGNALING | met | C7 | 1.662892555 | -0.148877487 | 3.48637E-06 |
| P6593 | INFLAMMATORY_RESPONSE | met | C7 | -2.089273109 | -0.185802121 | 1.79055E-03 |
| P6593 | KRAS_SIGNALING_DN | met | C7 | -2.647079973 | 0.120236713 | 2.39070E-11 |
| P6593 | KRAS_SIGNALING_DN | met | C1 | -2.647079973 | 0.302627928 | 2.39070E-11 |
| P6593 | KRAS_SIGNALING_DN | met | C8 | -2.647079973 | 0.25066592 | 1.39162E-09 |
| P6593 | KRAS_SIGNALING_DN | met | C5 | -2.647079973 | 0.230097659 | 2.39070E-11 |
| P6593 | KRAS_SIGNALING_DN | met | C3 | -2.647079973 | 0.353686858 | 2.39070E-11 |
| P6593 | KRAS_SIGNALING_DN | met | C4 | -2.647079973 | 0.428876877 | 2.39070E-11 |
| P6593 | KRAS_SIGNALING_DN | met | C6 | -2.647079973 | 0.495709839 | 2.39925E-11 |
| P6593 | KRAS_SIGNALING_DN | met | C2 | -2.647079973 | 0.465178179 | 2.45164E-11 |
| P6593 | KRAS_SIGNALING_UP | met | C7 | -2.424858777 | 0.50897544 | 2.39070E-11 |
| P6593 | KRAS_SIGNALING_UP | met | C1 | -2.424858777 | 1.192514917 | 2.42596E-11 |
| P6593 | KRAS_SIGNALING_UP | met | C5 | -2.424858777 | 0.123970029 | 8.69850E-03 |
| P6593 | KRAS_SIGNALING_UP | met | C3 | -2.424858777 | 0.490062831 | 1.11935E-03 |
| P6593 | KRAS_SIGNALING_UP | met | C4 | -2.424858777 | -0.247560979 | 2.69317E-02 |
| P6593 | SPERMATOGENESIS | met | C7 | -2.650218197 | 0.100847126 | 2.39070E-11 |
| P6593 | SPERMATOGENESIS | met | C1 | -2.650218197 | 0.303065343 | 2.39070E-11 |
| P6593 | SPERMATOGENESIS | met | C8 | -2.650218197 | 0.368965618 | 1.00549E-07 |
| P6593 | SPERMATOGENESIS | met | C5 | -2.650218197 | 0.373702601 | 2.39361E-11 |
| P6593 | SPERMATOGENESIS | met | C3 | -2.650218197 | 0.291035323 | 2.39794E-11 |

| | | | | | | |
|---|---|---|---|---|---|---|
| P6593 | SPERMATOGENESIS | met | C4 | -2.650218197 | 0.29156937 | 2.39080E-11 |
| P6593 | SPERMATOGENESIS | met | C6 | -2.650218197 | 0.508840936 | 2.39803E-11 |
| P6593 | SPERMATOGENESIS | met | C2 | -2.650218197 | 0.41219188 | 2.34141E-09 |
| P6593 | TNFA_SIGNALING_VIA_NFKB | met | C7 | 2.501013155 | 0.260962661 | 2.39070E-11 |
| P6593 | TNFA_SIGNALING_VIA_NFKB | met | C1 | 2.501013155 | -0.006892169 | 2.39945E-11 |
| P6593 | TNFA_SIGNALING_VIA_NFKB | met | C8 | 2.501013155 | -0.047373893 | 9.33849E-03 |
| P6593 | TNFA_SIGNALING_VIA_NFKB | met | C5 | 2.501013155 | -0.406674526 | 2.46162E-11 |
| P6593 | TNFA_SIGNALING_VIA_NFKB | met | C3 | 2.501013155 | -0.420974572 | 2.44545E-11 |
| P6593 | TNFA_SIGNALING_VIA_NFKB | met | C4 | 2.501013155 | -0.35600792 | 2.40589E-11 |
| P6593 | TNFA_SIGNALING_VIA_NFKB | met | C6 | 2.501013155 | -0.910056987 | 6.52722E-10 |
| P6593 | TNFA_SIGNALING_VIA_NFKB | met | C2 | 2.501013155 | -0.61399575 | 6.15341E-05 |
| P6593 | UNFOLDED_PROTEIN_RESPONSE | met | C7 | 2.658474332 | -0.161602409 | 2.39070E-11 |
| P6593 | UNFOLDED_PROTEIN_RESPONSE | met | C1 | 2.658474332 | -0.352598321 | 2.39070E-11 |
| P6593 | UNFOLDED_PROTEIN_RESPONSE | met | C8 | 2.658474332 | -0.284741656 | 2.40352E-11 |
| P6593 | UNFOLDED_PROTEIN_RESPONSE | met | C5 | 2.658474332 | -0.318059563 | 2.39070E-11 |
| P6593 | UNFOLDED_PROTEIN_RESPONSE | met | C3 | 2.658474332 | -0.334716798 | 2.39070E-11 |
| P6593 | UNFOLDED_PROTEIN_RESPONSE | met | C4 | 2.658474332 | -0.43237279 | 2.39070E-11 |
| P6593 | UNFOLDED_PROTEIN_RESPONSE | met | C6 | 2.658474332 | -0.40974906 | 2.39070E-11 |
| P6593 | UNFOLDED_PROTEIN_RESPONSE | met | C2 | 2.658474332 | -0.364633735 | 2.39962E-11 |
| P6593 | WNT_BETA_CATENIN_SIGNALING | met | C7 | -2.577591785 | -0.029842366 | 2.39070E-11 |
| P6593 | WNT_BETA_CATENIN_SIGNALING | met | C1 | -2.577591785 | 0.137062378 | 2.39078E-11 |
| P6593 | WNT_BETA_CATENIN_SIGNALING | met | C8 | -2.577591785 | 0.659467045 | 4.04212E-05 |
| P6593 | WNT_BETA_CATENIN_SIGNALING | met | C5 | -2.577591785 | 0.264534975 | 2.40791E-11 |
| P6593 | WNT_BETA_CATENIN_SIGNALING | met | C3 | -2.577591785 | 0.306091443 | 2.40409E-11 |
| P6593 | WNT_BETA_CATENIN_SIGNALING | met | C4 | -2.577591785 | 0.239089397 | 2.39965E-11 |
| P6593 | WNT_BETA_CATENIN_SIGNALING | met | C6 | -2.577591785 | 0.800113288 | 7.89250E-11 |
| P6593 | WNT_BETA_CATENIN_SIGNALING | met | C2 | -2.577591785 | 0.201075624 | 2.02045E-04 |
